# The impact of sex, age, and genetic ancestry on DNA methylation across tissues

**DOI:** 10.1101/2025.04.29.651179

**Authors:** Winona Oliveros, Jose Miguel Ramirez, Marta Melé

## Abstract

Understanding the consequences of individual DNA methylation variation is crucial for advancing our knowledge of human biology and disease. Yet, the collective impact of individual traits on DNA methylation and their downstream effects on gene expression across human tissues remain poorly understood. Here, we quantify the contributions of sex, age, genetic ancestry, and BMI on autosomal DNA methylation variation across 9 human tissues and 424 individuals from the Genotype-Tissue Expression project. We show that genetic ancestry and age have a greater impact on DNA methylation than sex, with aging effects being more widespread but less pronounced. On average, less than 10% of the gene expression variation in sex, age, and ancestry are mediated by DNA methylation differences, with ancestry showing the largest proportion of mediation. We further show that ancestry-associated DNA methylation differences accumulate at CpG sites with extreme methylation states and are largely under genetic control. The female autosomal genome exhibits consistent hypermethylation across tissues at Polycomb-repressed regions. Ultimately, we show that age-related Polycomb target hypermethylation is observed across multiple tissues, but not in the gonads. Our multi-individual, multi-tissue approach defines the key drivers of human DNA methylation variation in healthy conditions, establishing a baseline for the interpretation of DNA methylation changes in disease contexts.

## Introduction

The deposition and maintenance of DNA methylation is essential for normal mammalian development. DNA methylation occurs when a methyl group (-CH3) is transferred into, almost exclusively, the C5 position of the cytosine in CpG dinucleotides(Jin et al. 2011; Jones and Taylor 1980; 2009; Smith and Meissner 2013). DNA methylation is fundamental for the regulation of X-inactivation(Smith and Meissner 2013), genomic imprinting(Jones 2012; Robertson 2005), and the repression of repetitive elements(Robertson 2005). Most (60–80%) of the 28 million CpGs in the human genome are methylated, which has mainly been associated with gene silencing(Jones 2012). The remaining 20–40% of unmethylated CpGs are mostly located in CpG islands, which are genomic regions with very high CpG content that are resistant to DNA methylation. However, the role of DNA methylation on transcriptional regulation depends on the genomic context(Jones 1999). DNA methylation at promoters, which are usually located in genomic regions containing CpG islands, is associated with long-term gene silencing(Babenko et al. 2017; Jones 2012), whereas DNA methylation at the gene body can, in certain contexts, stimulate transcription elongation(Lister et al. 2009; Jones 2012). Importantly, aberrations in DNA methylation are associated with many diseases(Robertson 2005), such as various cancer types(Widschwendter et al. 2004; Gama-Sosa et al. 1983; Adorján et al. 2002; De Smet et al. 1996), autoimmune and inflammatory diseases(Paul et al. 2016; Tahara et al. 2017; Hall et al. 2013; Yang et al. 2012; Liu et al. 2013), and cardiovascular diseases(Dekkers et al. 2016; Watson et al. 2016; Zaina et al. 2014). Thus, DNA methylation plays a fundamental role in genome regulation.

DNA methylation can vary between individuals, influenced by both intrinsic (e.g., age, genetics, and sex), and environmental factors. Age-related changes in DNA methylation are among the most extensively studied, including many individual CpGs for which methylation status has been reported to correlate with age(Horvath and Raj 2018). In particular, it has been shown that with age, DNA becomes more hypermethylated at bivalent chromatin regions and CpG islands(Rakyan et al. 2010; Yuan et al. 2015), whereas it becomes more hypomethylated at open-sea regions (i.e., isolated CpGs)(Lu et al. 2023; Christensen et al. 2009). In fact, the accumulation of DNA methylation changes can accurately predict chronological age and mortality(Horvath 2013; Beynon et al. 2022; McCrory et al. 2020). Sex-based differences in DNA methylation further contribute to the complexity of inter-individual variation. Methylation profiles exhibit pronounced differences between males and females across various tissues, including both sex chromosomes and autosomes(Davegårdh et al. 2019; Liu et al. 2010; Singmann et al. 2015). For example, female genomes display higher levels of autosomal methylation than male genomes do in whole blood(Grant et al. 2022). Finally, genetic variation significantly influences DNA methylation through mechanisms such as methylation quantitative trait loci (meQTLs), which link specific genetic variants to methylation levels at particular CpG sites. This genetic influence is evident in the differential methylation patterns observed across human populations in whole blood(Husquin et al. 2018; Galanter et al. 2017; Giri et al. 2017; Bell et al. 2011), which reflect both the ancestral genetic backgrounds and the impact of environmental exposures.

Despite the valuable insights provided by these studies, the study of DNA methylation variation has predominantly been confined to a limited number of tissues and single traits(Loyfer et al. 2023; Husquin et al. 2018; Yuan et al. 2015), with very limited efforts trying to disentangle its regulatory effect(Dozmorov 2015; Hannum et al. 2013). A recent study in whole blood dissected the collective contribution of different environmental and intrinsic factors to DNA methylation variation(Bergstedt et al. 2022). This study revealed that many factors, including age and sex, have a direct effect on DNA methylation, even when controlling for cell type composition effects. However, beyond whole blood, the comparative impact of individual traits, such as age, genetic ancestry, sex, and body mass index (BMI), across tissues on DNA methylation and its effect on transcriptional regulation, remains largely unexplored.

In this study, we leveraged Genotype-Tissue Expression (GTEx) data to systematically investigate the relationships among sex, age, ancestry, and BMI, with DNA methylation variation across 9 human tissues encompassing 856 samples from a total of 424 individuals. We identified differentially methylated positions (DMPs; capturing differences in mean methylation values) and differentially variably methylated positions (DVPs; capturing differences in variance of methylation) across tissues with different individual traits and their associations with gene expression. We identified common, trait-and tissue-specific patterns, which we further characterized via publicly available data, such as chromatin states, to determine their functional consequences. Overall, our multi-trait and multi-tissue analysis provides an exhaustive characterization of the global and tissue-specific impact of age, sex, ancestry, and BMI on DNA methylation variation.

## RESULTS

### Individually variable methylated CpGs are preferentially located at promoters

DNA methylation is known to vary both between individuals and between tissues(Yuan et al. 2015; Husquin et al. 2018; Davegårdh et al. 2019). However, the extent to which DNA methylation varies more across tissues than across individuals has only been estimated with few donors(Byun et al. 2009; Loyfer et al. 2023). In addition, whether DNA methylation variability is comparable to gene expression variability remains unexplored. To first quantify how DNA methylation varies across individuals and tissues, we used data from 9 GTEx tissues from 424 individuals across autosomal 754,054 CpG sites from the Illumina EPIC methylation array (**Fig. 1A, Fig. S1A**)(Oliva et al. 2023). We used a variance partitioning approach to quantify the contribution of individual and tissue to DNA methylation variation, controlling for known sources of technical variation and other confounders, such as cell-type composition (see Methods). Variation in DNA methylation is substantially greater among tissues (mean ∼53% of the total variance in DNA methylation) than among individuals (mean ∼3.7% of the total variance) (**Fig. 1B**), as previously reported(Byun et al. 2009; Loyfer et al. 2023). We then calculated the proportion of gene expression variation explained by individual and tissue using the same tissues and approach as with DNA methylation. Compared with methylation, gene expression varies less between tissues (mean ∼45% of the total variance, Mann‒Whitney‒Wilcoxon test p value < 2.2e‒16, **Fig. 1C**) and individuals (mean ∼2.6% of the total variance, Mann‒Whitney‒Wilcoxon test p value = 5.815e‒05), which is consistent with previous studies with smaller sample sizes(Melé et al. 2015). Thus, tissue and individual factors explain a larger proportion of DNA methylation variability than gene expression variability does, probably reflecting the higher stability of DNA methylation and noisier nature of RNA assays.

We then evaluated whether DNA methylation variation correlates with gene expression variation. To do so, we defined two sets of highly variable genes (>50% variance explained either by tissue or individual for DNA methylation – using the EPIC annotation – or gene expression, respectively) and looked for overlaps. Although there was no overlap between individually variable genes at the DNA methylation (2393 genes) and gene expression (11 genes) levels, we found significantly higher gene expression variance between individuals for genes that we detected to be highly individually variable in DNA methylation (Mann‒Whitney‒Wilcoxon test p value < 0.05) (**Fig. S1B‒C**). Accordingly, we found the same pattern for tissue variable genes (21146 and 7452 tissue-variable genes at DNA methylation and gene expression levels, respectively, **Fig. S1D**). This finding indicates that DNA methylation, although significant, might have a modest contribution to explaining inter-individual and inter-tissue variation in gene expression.

Next, we wanted to explore whether the proportion of DNA methylation variation explained by tissue or individual variation changes depending on the genomic context of the CpGs. We divided the CpGs into enhancers, promoters, gene bodies, or intergenic regions according to the EPIC array annotation and found that enhancer-associated CpGs have higher tissue variability than the rest (OR= 3.81, FDR= 0, two-tailed Fisher’s exact test), with promoter-associated CpGs being highly stable across tissues (**Fig. S1E-F)**. This observation is consistent with previous studies showing enrichment of cell type-specific methylation patterns at enhancer regions(Dor and Cedar 2018; Loyfer et al. 2023). Although the influence of the genomic context on the proportion of variation explained by individual is modest (**Fig. S1E**), we detected a significantly larger impact at promoters (OR= 1.55, FDR= 2.3e-29, two-tailed Fisher’s exact test; **Fig. 1D**). We then evaluated whether the contribution of DNA methylation variation to gene expression variation changes on the basis of the genomic context(Jones 2012). When stratifying the inter-individual gene expression variance between genes highly variable in DNA methylation for each genomic location, we found that only DNA methylation variation in promoters significantly correlates with inter-individual gene expression variation (**Fig. 1E, Fig. S1G-H**).

Finally, we wanted to explore whether highly tissue or highly individually variable CpGs were associated with genes of particular functions by functional enrichment analysis. Highly tissue-variable CpGs are more often located near genes involved in developmental and metabolic processes (functional enrichment, FDR < 0.05) (**Table S1**), which likely reflects the developmental fine-tuning of tissue-specific gene expression and is consistent with previous observations where cell types cluster on the basis of developmental origin(Loyfer et al. 2023). Highly individually variable CpGs (**4810** CpGs) are enriched in genes belonging to antigen processing and presentation pathways (functional enrichment, FDR < 0.05; **Table S1; Fig. 1F**). This enrichment is driven mostly by the presence of highly inter-individual variable CpGs located in the HLA genes located in the MHC locus (**Table S1**), which is one of the most polymorphic loci in vertebrates(Vogel et al. 1999) and has been suggested to be transcriptionally regulated by DNA methylation(Rakyan et al. 2004). Consistent with this observation, genes belonging to antigen processing and presentation pathways have higher inter-individual gene expression variation than other genes (Mann-Whitney-Wilcoxon Test p.value = 0.0012) (**Fig. 1G**).

Overall, we find higher tissue and individual variability of DNA methylation compared to gene expression. Interestingly, highly individually variable CpGs are located in promoters and enriched in the HLA genes, moderately correlating with their gene expression variability and potentially reflecting differences in immune responses between individuals.

**Figure 1.**
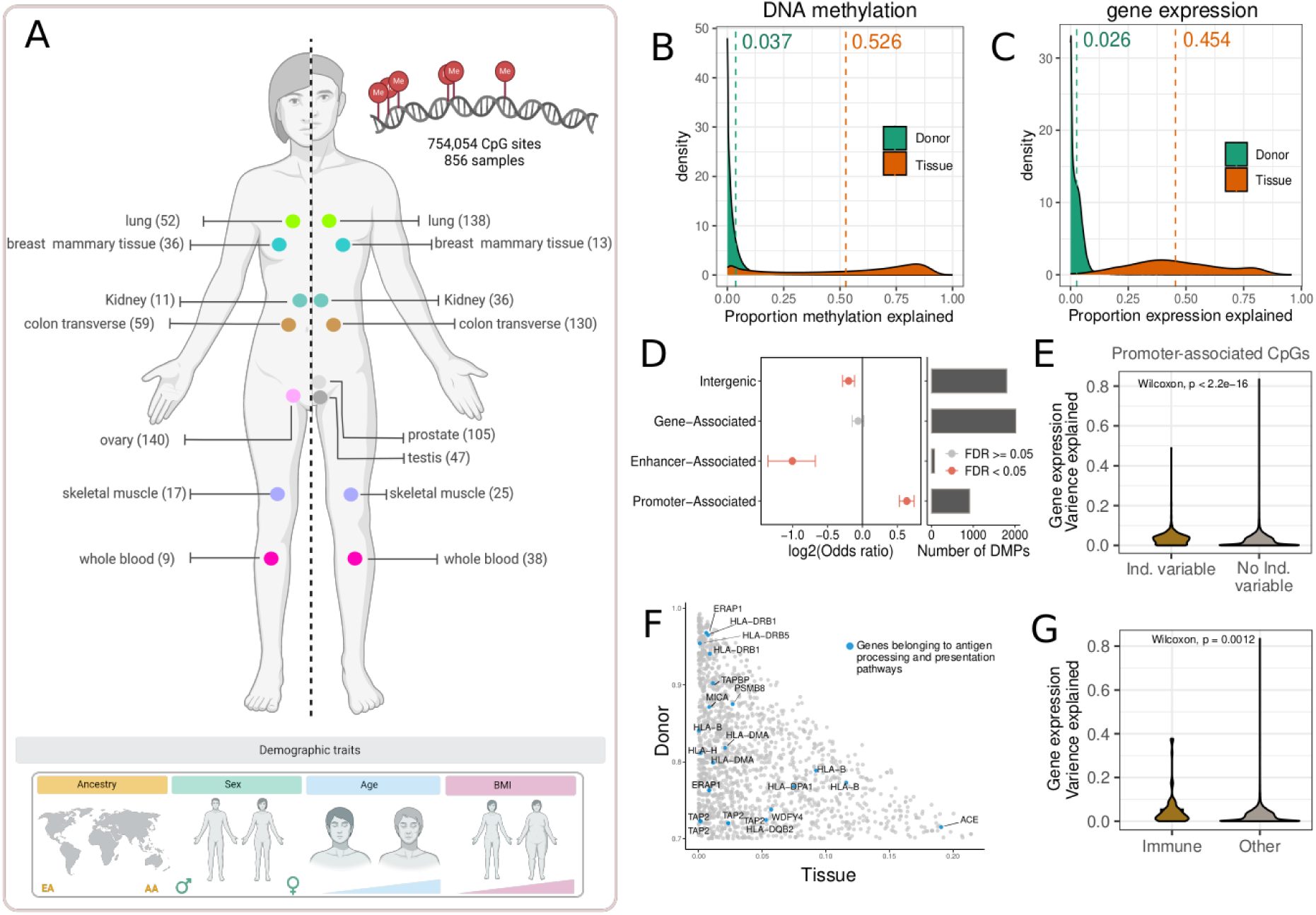
DNA methylation variability across individuals significantly affects promoters. **A.** Overview of the DNA methylation data, with the number of female samples (left) and the number of male samples (right) for each tissue. The number of analyzed CpGs and the total number of samples are shown. **B**. Proportion of DNA methylation variation explained by individual (green) and tissue (orange). The mean numbers are shown. **C.** Proportion of gene expression variation explained by individual (green) and tissue (orange). The mean numbers are shown. **D. Left.** Enrichment of each genomic location on individually variable CpGs (>50% variance explained either by individual for DNA methylation). Dots are colored according to the significance of the enrichment (two-tailed Fisher’s exact test). The confidence interval is shown. **Right.** Number of individually variable CpGs within each genomic location. The total numbers of CpGs tested in promoters, enhancers, and gene bodies are 108,810, 25,287, and 370,343. **E.** Proportion of gene expression variation explained for genes that are individually-variable (brown) or not individually-variable (gray) at the DNA methylation level of their promoters. p values from Mann‒Whitney‒Wilcoxon test. **F.** Zoom-in of Individually variable CpGs. The X-axis represents the proportion of DNA methylation variation explained by tissue. The Y-axis represents the proportion of DNA methylation variation explained by individual. The blue dots represent CpGs associated with genes belonging to antigen processing and presentation pathways. **G.** Proportion of gene expression variation explained for genes belonging to antigen processing and presentation pathways (blue) versus other genes (gray). p values from Mann‒Whitney‒Wilcoxon test.

### Age and ancestry have a larger impact on DNA methylation than sex across tissues

Genetic effects, age, and sex simultaneously contribute to DNA methylation variation in whole blood, with genetic effects having the strongest impact(Bergstedt et al. 2022). To further study the simultaneous association between individual traits and DNA methylation variation beyond whole blood, we quantified DNA methylation changes with genetic ancestry, sex, age, and BMI simultaneously across 9 human tissues. We identified differentially methylated positions (DMPs, adjusted p value < 0.05) for each individual trait per tissue using linear fixed-effects models controlling for known sources of technical variation and other confounders, such as cell-type composition (see Methods). In most tissues, age has the largest number of DMPs, followed by ancestry and sex (**Fig. 2A**). However, tissues with smaller sample sizes have more ancestry-DMPs than age-DMPs (**Fig. 2A**). This is likely because ancestry-DMPs have larger effect sizes (effect sizes of ancestry-DMPs vs. age-DMPs, Mann‒Whitney‒Wilcoxon test p value < 2.2e‒16 in all tissues) (**Fig. S2A**) and thus can be detected in tissues with low sample sizes. BMI shows no association with DNA methylation changes in the studied tissues, consistent with previous observations in whole blood(Wahl et al. 2017; Dick et al. 2014). The largest number of age-DMPs and ancestry-DMPs are located in the colon and lung respectively, which have the largest sample sizes. Conversely, skeletal muscle, despite not having many samples, has the largest number of sex-DMPs, consistent with previous reports of widespread sex DNA methylation and physiological differences(Maher et al. 2009; Landen et al. 2019, 2021). Downsampling tissues to the same number of samples shows that skeletal muscle remains the tissue with the highest number of sex-DMPs and that ancestry has the largest number of DMPs across tissues (**Fig. S2B**).

Next, we tested the specific contribution of each individual trait to DNA methylation variation per tissue using a variance partitioning approach (see Methods). We found that in every tissue, the individual trait with the largest number of DMPs also explains most of the variation in DNA methylation variation (**Fig. 2A**). In general, however, the percentage of total variation explained by individual traits per CpG is low (mean DNA methylation variation explained: 10.7% ancestry, 6% age, 3.8% sex, and 0.9% BMI). From the 23,120 DMPs for which a single trait explained more than 10% of their inter-individual DNA methylation variation in at least one tissue, age had the largest number of DMPs, followed by ancestry and sex (age 43%, ancestry 42%, sex 15%, examples of CpGs with a large proportion of DNA methylation variation explained by an individual trait in **Fig. 2B**). Among those, some are known to be related to trait-associated phenotypes, such as cg01878807 and cg05171021, two CpGs located on the promoter of the *DHRS4* gene (ADME gene, highly related to the absorption, distribution, metabolism, and excretion of drugs), where ancestry explains more than 10% of the variability in DNA methylation. Higher DNA methylation of both CpGs in African-American individuals is correlated with lower gene expression(Chu and Yang 2017) and could impact ethnic differences in treatment response and drug ADME(Ramos et al. 2014).

Finally, we evaluated whether the contribution of each individual trait to DNA methylation variation in each tissue could be correlated with the contribution to gene expression variation. To compare both data modalities, we studied the genes associated with DNA methylation changes in the same tissues (see Methods). Compared to methylation, the percentage of total gene expression variation explained by each individual trait per gene is lower (mean gene expression variation explained: 1.3% ancestry, 2.7% age, 1.7% sex, and 1.9% BMI). Importantly, the contribution of each individual trait to explaining gene expression variation mirrors previous results(García-Pérez et al. 2022) but is different from that of DNA methylation, with an overall higher contribution from BMI and a lower contribution from ancestry (**Fig. S2C**).

Overall, age and ancestry have larger contributions to inter-individual variation in DNA methylation variation than sex, with aging effects being more abundant but weaker. Interestingly, we report a higher contribution of ancestry, age, and sex to DNA methylation variation than to gene expression, probably reflecting the higher stability of DNA methylation and noisier nature of RNA assays.

### Few age-, sex-, or ancestry-related expression differences are mediated by DNA methylation

DNA methylation is a key regulatory layer of the genome, but to what extent methylation differences influence, or reflect, changes in gene expression across individuals remains unexplored. To further explore their putative correlation, we asked if previously observed differences in gene expression variation with age, sex, and ancestry(García-Pérez et al. 2022) could be explained by changes in DNA methylation. To address this, we first selected the DMPs that had an associated differentially expressed gene (DEG) for the same trait and tissue combination in a previous study(García-Pérez et al. 2022) (see Methods). The proportion of DMP-DEG pairs was very low (0.4%-13.7%), suggesting that most expression differences with respect to age, sex, or ancestry might not be correlated with methylation differences. In fact, fewer than 10% of the gene expression differences correlated with DNA methylation changes in the DMP-DEG pairs (**Fig. 2C, Fig. S2D-E**). Ancestry was the individual trait with the highest percentage of correlated genes (mean ∼8%, maximum 20% of correlated DEGs). Contrarily, age showed overall few significant correlations (mean 4% correlated DEGs), consistent with previous results(Yuan et al. 2015). The exception to this trend is ovary, which yielded 432 correlated DMP-DEGs (corresponding to 239 DEGs, 10%) located near genes involved in blood circulation pathways (**Table S2**), probably reflecting the reorganization of the tissue during menopause. As expected, most correlations involving CpGs located at promoters are negative compared with a lower proportion of negative correlations in enhancers and gene bodies (**Fig. 2D, Table S3**). We explored whether the low correlation between gene expression and DNA methylation could be explained by the limited statistical power of our differential DNA methylation analysis, which includes fewer samples than the previously published differential gene expression analysis(García-Pérez et al. 2022). To address this, we correlated the significant trait-DMPs (p value nominal < 0.05) with the corresponding DEGs (see Methods). The proportion of gene expression differences correlated with DNA methylation substantially increased (up to 44% correlated DEGs in ovaries with genetic ancestry) (**Fig S2F**), suggesting that other genes transcriptionally correlated with DNA methylation could be missing from existing correlation analyses.

Finally, we further explored the putative causal relationship between DNA methylation and gene expression by testing whether the effect of each individual trait on gene expression could be mediated by changes in DNA methylation. To address this, we applied a regularized mediation analysis(Schaid and Sinnwell 2020) to estimate the average causal mediation effect of DNA methylation on every DEG per trait and tissue (see Methods). On average, ∼4% of the DEGs are mediated by DNA methylation (**Fig. 2E, Table S4**). In this mediation analysis, we included all the DMPs associated with a DEG in the same model as multiple mediators. Therefore, the similar proportion of mediated effects (mean ∼4%) compared with our previous correlation analysis (mean ∼10%) probably indicates that when there is a putative mediation effect of DNA methylation on gene expression, a single CpG is sufficient to detect it (**Fig. S2G, Table S4**).

Together, these results indicate that age-, sex- and ancestry-related DNA methylation changes in single CpGs do not normally mediate gene expression changes under healthy conditions, probably reflecting the noisier nature of RNA compared to more stable changes in DNA methylation(Furukawa et al. 2016; Raj and van Oudenaarden 2008).

### DNA methylation variability is higher in individuals of African descent

DNA methylation has been shown to increase in variability with age across tissues(Slieker et al. 2016; Bergstedt et al. 2022). We thus wanted to test whether sex and ancestry could also be associated with increased DNA methylation variability and compare their effects to age. We identified differential variable positions (DVPs, adjusted p value < 0.05) for age, sex, and ancestry using a linear fixed-effect model controlling for the same confounders as in our previous differential DNA methylation analysis (see Methods). In all tissues, ancestry has the largest number of DVPs (**Fig. 2F**). Interestingly, ∼99% of the ancestry-DVPs show larger variability in individuals of African descent (binomial test; FDR < 0.05 in all tissues, **Fig. 2G**), consistent with African individuals being more genetically diverse(2015a). We identified a limited number of age- and sex-related DVPs. Consistent with this, the overlap between DMPs and DVPs is significant in all tissues only for ancestry (∼25% of DMPs are also DVPs, two-tailed Fisher’s exact test, FDR < 0.05), likely because both capture genetic effects and have larger effect sizes (**Fig. S2A, Fig. S2H**) compared to age and sex.

We then examined whether changes in methylation variability with sex, age, or ancestry could have an association with gene expression. To address this, we first applied the same methodology to detect differentially variable genes (DVGs) using the gene expression data available for the same tissues from the GTEx dataset. On average, we detect ∼290 age-DVGs, ∼1800 sex-DVGs and ∼1020 ancestry-DVGs across tissues (**Fig. S2I)**. In 99% of cases, ancestry-DVGs increase in expression variability in African individuals (binomial test; FDR < 0.05 in all tissues except testes, **Fig. 2H**). However, we find almost no overlap between trait-DVP-DVGs (mean overlap: 0 age-DVP-DVG, 1 sex-DVP-DVG, and 11 ancestry-DVP-DVG, **Fig. S2J**). Our results suggest that variable CpGs have little effect on the regulation of gene expression.

Overall, we report widespread differences in DNA methylation between populations, which might reflect the higher genetic variability found in African populations and/or environmental effects.

**Figure 2.**
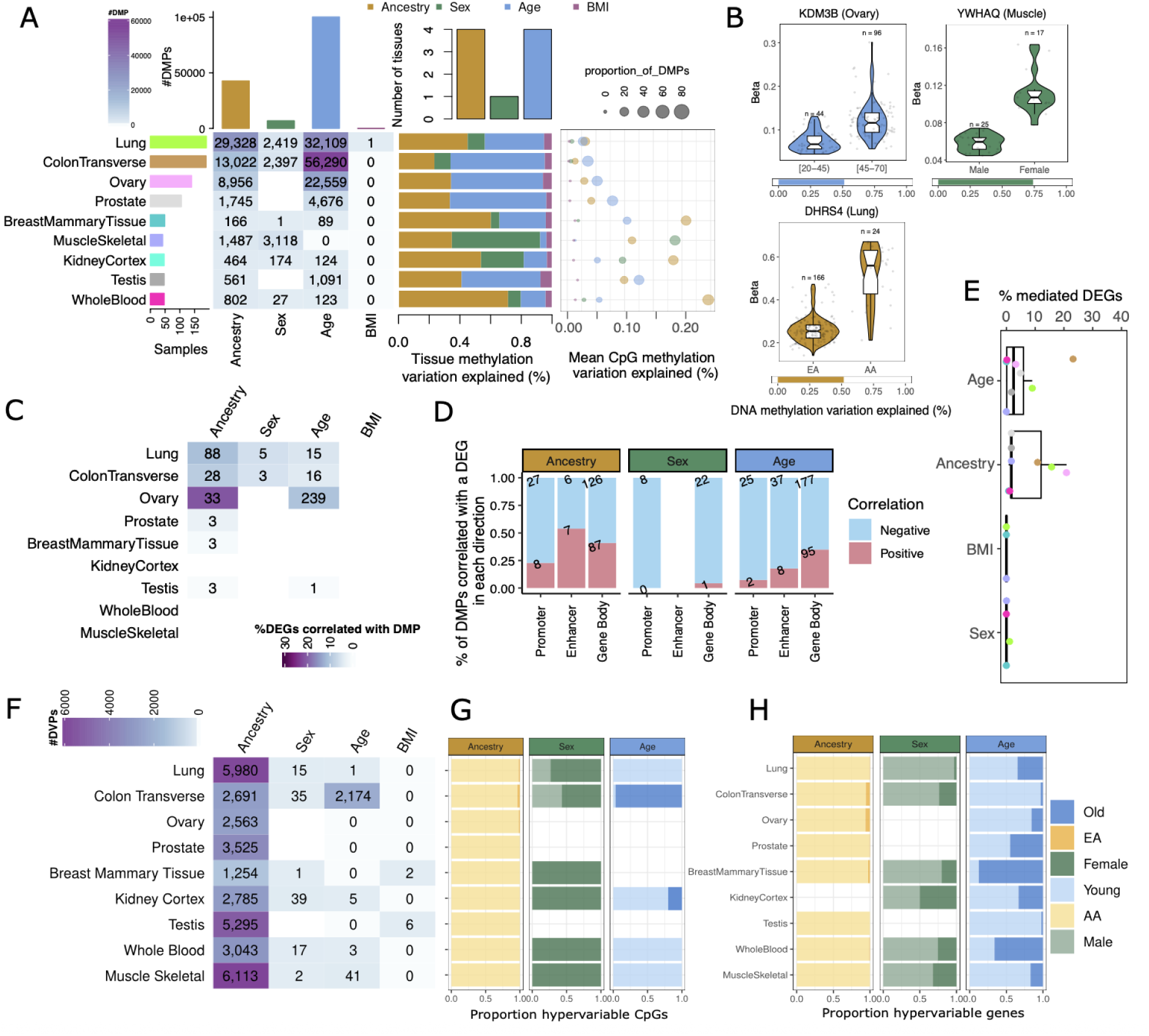
Age and ancestry significantly contribute to DNA methylation variation, with limited association with gene expression changes. **A. (Left)** Number of DMPs (adjusted p value < 0.05) per tissue and individual trait. Top bars show the number of CpGs differentially methylated by each individual trait. Left bars represent the sample size. **(Center)** Proportion of total tissue DNA methylation variation explained by each individual trait. Top bars show the number of tissues for which each individual trait explains the largest proportion of variation. **(Right)** Mean DNA methylation variation explained by each individual trait in each tissue. **B.** Examples of CpGs with a large proportion of DNA methylation variation explained by an individual trait. The aging variable was categorized for visualization purposes. Top violin plots represent the beta values of each CpG across individuals, and the bottom bars represent the proportion of DNA methylation variation explained by each individual trait. **C.** Number of DEGs per tissue and individual trait correlated with at least 1 DMP. Cells are colored according to the proportion of DEGs correlated with DMPs out of the total number of DEGs with a CpG. Tissues are sorted by sample size. Refer to A for sample size information. **D.** Proportion and number of DMPs correlated with DEGs per genomic location and individual trait, colored according to the direction of correlation. **E.** Percentage of DEGs per tissue and individual trait that are mediated by DNA methylation. Dots are colored according to tissue; see barplot in panel A. **F.** Number of DVPs (adjusted p value < 0.05) per tissue and individual trait. Refer to A for sample size information. Cells are colored according to the number of DVPs. **G.** Proportion of hyper-variably methylated CpGs per tissue and individual trait. **H.** Proportion of hyper-variably expressed genes per tissue and individual trait. Old/EA/Female refer to hyper-variable CpGs/genes in the model; Young/AA/Male refer to hypo-variable CpGs/genes in the model. For female/male sample size, refer to Figure 1A. Age and Ancestry are modeled as continuous variables (see Methods).

### Africans and Europeans show distinct methylation patterns at TSSs and quiescent regions

To better understand the relationship between genetic ancestry and epigenetics, we tested whether ancestry-DMPs are located in regions of the genome with specific chromatin states across tissues (see Methods). We find that hypermethylated ancestry-DMPs in European-descent individuals are more often located in quiescent regions whereas ancestry-DMPs hypermethylated in African-descent individuals are consistently enriched in TSSs (two-tailed Fisher’s exact test, FDR < 0.05) (**Fig. S3A**) across tissues. When selecting the shared ancestry-DMPs in the same direction (hypermethylated in 3 or more tissues), we observe the same enrichment (two-tailed Fisher’s exact test, FDR < 0.05). (**Fig. 3A**). Interestingly, quiescent regions are, on average, the chromatin states with the highest methylation levels, whereas promoters have the lowest methylation levels (**Fig. S3B**). Hence, we evaluated whether our previous observation was driven by Africans being more diverse; and thus, regions with extreme DNA methylation profiles, such as promoters and quiescent regions, fluctuate in only one direction. To test this, we selected CpGs with extreme methylation values (i.e., CpGs with average methylation levels > 0.9 and < 0.1) and tested whether ancestry-DMPs are significantly enriched in these extremely methylated and unmethylated CpGs. Indeed, we found that, on average, CpGs with high DNA methylation levels were hypomethylated in Africans compared with Europeans (**Fig 3B**), and, on average, CpGs with low DNA methylation levels were hypermethylated in Africans (**Fig S3C**). This observation is consistent with the higher genetic variability observed in Africans(2015a) and the larger DNA methylation variability we observed in African-descent individuals (**Fig. 2G**).

We then hypothesized that the enrichment in TSSs of ancestry-DMPs hypermethylated in Africans (**Fig. 3A, S3A**) might have an association with the expression of the associated genes, as higher methylation at promoter regions is usually associated with lower gene expression(Jones 1999). To test this hypothesis, we compared the mean expression levels of genes with hypermethylated ancestry-DMPs in their promoters in Africans. As expected, we found that the expression of these genes is lower in Africans than in Europeans (Wilcoxon paired test; p value<0.05).

Overall, our results show that widespread differences in DNA methylation profiles between African and European populations localize to regions of the genome with extreme methylation profiles (either completely methylated or unmethylated). This pattern likely reflects the higher genetic diversity and DNA methylation variability observed in African individuals, as variability is more easily detected in such regions where methylation values can shift in only one direction. Finally, we show that DNA methylation variation in promoter regions has a small, but significantly consistent, influence on the gene expression of associated genes.

### Genetic effects explain ∼70% of the differences in DNA methylation between human populations across tissues

Ancestry-associated DNA methylation changes can reflect the impact of both genetic and environmental factors(Bell et al. 2011; Feil and Fraga 2012). A previous study in whole blood comparing methylation levels between a balanced number of European-descent and African-descent individuals estimated that 70% of ancestry-DMPs were driven by cis genetic effects(Husquin et al. 2018). Here, we wanted to quantify the proportion of ancestry-DMPs that are driven by genetic effects across tissues. To do this, we selected the list of *cis*-methylation quantitative trait loci (*cis*-mQTLs) (i.e., local associations between DNA methylation variation at CpGs and SNPs located within a 500-kb window) identified in the same samples in a previous study(Oliva et al. 2023) and overlapped them with the ancestry-DMPs in the same tissue. On average, we find 25 to 50% of ancestry-DMPs in CpGs with associated *cis*-mQTLs (**Fig. 3C**). In the tissues with low sample sizes (whole blood, testis, kidney cortex, muscle, and breast), few of the ancestry-DMPs were previously associated with a *cis*-mQTL, suggesting that there might be genetic effects that have yet to be identified. However, we find that ancestry-related DMPs are significantly enriched in *cis*-mQTLs in 5 tissues (mCpGs) (two-tailed Fisher’s exact test, FDR < 0.05) (**Fig. S3D**). To test whether the observed differences in DNA methylation between African and European populations are explained by genetic effects, we included the associated *cis*-mQTLs in our differential methylation models and tested whether those CpGs were still differentially methylated using F tests (see Methods). We find that, on average, ∼70% of ancestry-DMPs that have an associated *cis*-mQTL can be explained by *cis-*genetic effects (here referred to as *cis*-driven DMPs) (**Fig. 3D**).

We then assessed whether the ancestry-related DVPs could also be driven by *cis*-genetic effects, as they are highly shared across tissues (**Fig. S3E**) and have a strong bias toward increased DNA methylation variability in African-descent individuals (**Fig. 2G**). Similar to DMPs, few DVPs are associated with *cis*-mQTLs (**Fig. 3E**). Nevertheless, when they are present, most (average of ∼75%) are explained by genetic variants (here referred to as *cis*-driven DVPs) (**Fig. 3F**).

Among the four analyzed individual traits, ancestry demonstrates the highest cross-tissue DMP sharing, with 25% (moderate sharing, 9419 CpGs) of its DMPs found in more than one tissue and 2% (high sharing, 903 CpGs) in 4 or more tissues (**Fig. 3G**). As expected, *cis*-driven DMPs were significantly more tissue-shared than DMPs without an identified *cis*-mQTL and more significantly shared than non-*cis*-driven DMPs (Mann-Whitney-Wilcoxon Test p value < 0.05; **Fig. 3H**). Additionally, non-*cis*-driven DMPs were more significantly shared than DMPs without an identified mQTL (Mann‒Whitney-Wilcoxon Test p value < 0.05, **Fig. 3H**), consistent with the observation that *cis-*mQTL effects are highly shared across tissues(Oliva et al. 2023).

*Cis*-driven DMPs were enriched in enhancers (**Fig. S3F**) consistent with the fact that *cis*-mQTLs are usually located in non-genic regulatory regions(Oliva et al. 2023; Husquin et al. 2018). As recent work has shown that chromatin states, specifically enhancers, might differ between African and European individuals(Aracena et al. 2024), we explored whether ancestry-DMPs could be located in regions that had different chromatin states between these human populations. To address this, we used PBMC chromHMM states for an African-descent and a European-descent individual and assayed the preferential location of ancestry-DMPs in each individual (see Methods). Overall, 38% of whole-blood ancestry-related DMPs were located in differently annotated genomic regions between African and European populations (OR = 1.23, p value = 0.0044; **Fig. S3G**), with higher divergence at regulatory and repressed regions (**Fig. 3I**). Indeed, DNA hypermethylation in European-descent individuals was preferentially located in European-specific enhancers (OR=1.94, FDR=0.004; **Fig. S3H**), whereas DNA hypermethylation in African-descent individuals was preferentially located in African-specific bivalent enhancers (OR=2.23, FDR=0.04) (**Fig. S3I)**.

We next sought to characterize the functional pathways altered by ancestry-related DNA methylation changes. No pathway was significantly altered by overall ancestry-DMPs; however, enhancer-associated ancestry-DMPs were enriched in wound-healing and inflammation-related processes in the lung and colon (**Table S5**). Wound healing has been reported to play a role in inflammation(2017; Koh and DiPietro; Mosser and Edwards 2008) and inflammatory diseases(Chen et al. 2016). Interestingly, 15 DM genes associated with wound healing are differentially expressed between populations during infection (8 DEGs in monocytes, 9 DEGs in CD8+ T cells, 8 DEGs in CD4+ T cells, 9 DEGs in NK cells and 11 DEGs in B cells), with 4 additional DM genes being differentially responsive during infection (*ABHD2*, *MAX*, *MYH9*, and *AHNAK* in monocytes and *AHNAK* in B cells)(Randolph et al. 2021). Additionally, one of the most tissue-shared ancestry-DMP, cg16999677 (**Fig. 3J**), is hypermethylated in European-descent individuals across 8 tissues, and is the CpG with its DNA methylation variation more strongly explained by genetic ancestry. This CpG is located in the gene body of the *ZDHHC11* gene, which plays a key role in the antiviral innate immune response(Liu et al. 2018), and has been recently identified as a positive modulator of *NF-κB* signaling, a critical transcription factor (TF) for inflammatory responses(Liu et al. 2021). Additionally, this gene was reported to be significantly overexpressed in African-American individuals in four tissues (two brain regions, the liver, and the thyroid)(García-Pérez et al. 2022), consistent with higher methylation of this gene in European-American individuals resulting in lower expression.

Our results suggest that although *cis-*genetic effects mainly drive systemic DNA methylation differences between African and European human populations, many potential *cis*-mQTLs have yet to be identified. Additionally, differences in DNA methylation between these populations significantly affect population-specific regulatory chromatin regions.

**Figure 3.**
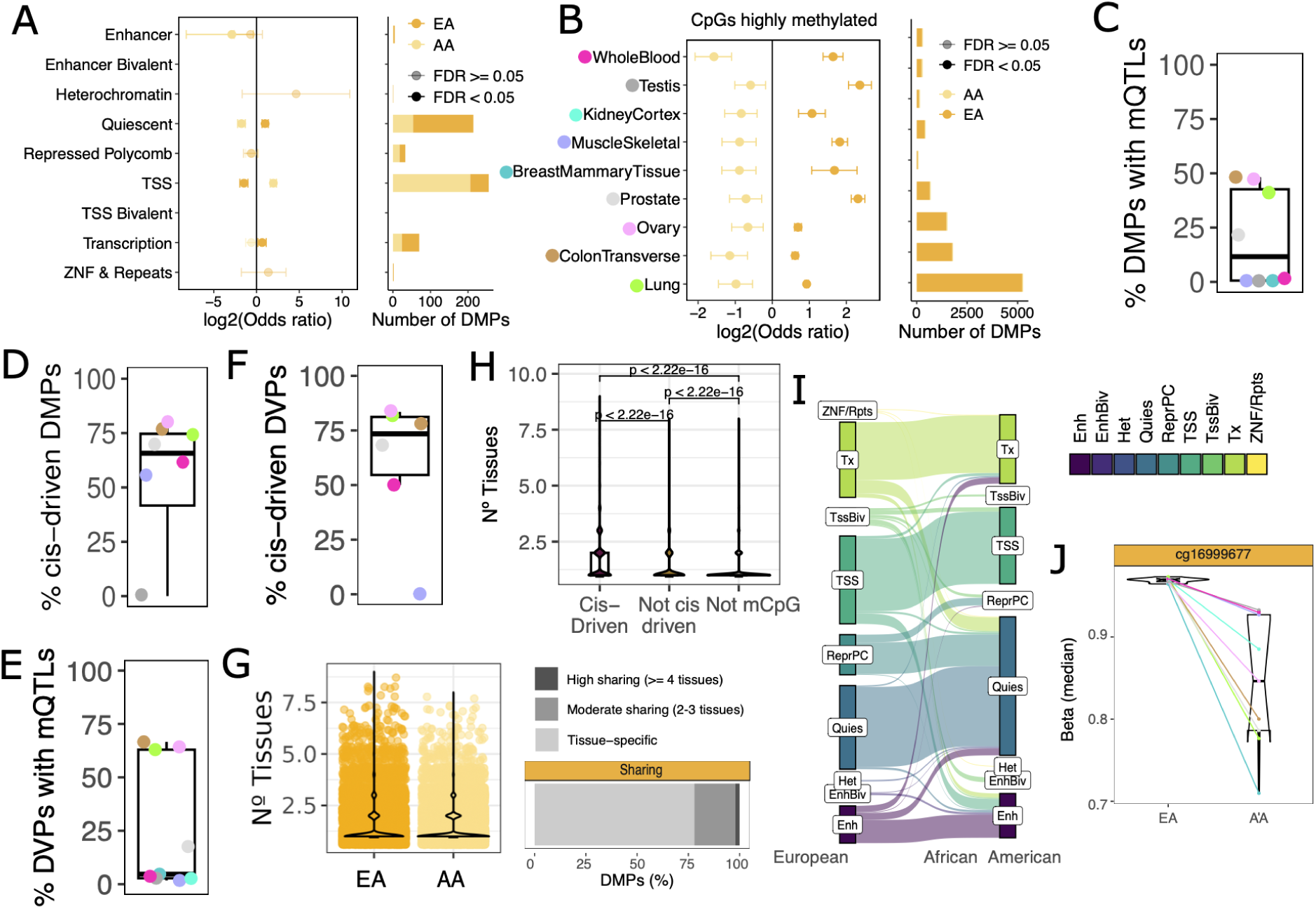
Ancestry-associated DNA methylation changes reflect chromatin context and are largely explained by genetic variation. **A.** Enrichment of shared ancestry-DMPs (CpGs differentially methylated with ancestry across 3 or more tissues) in chromatin states. Left. Enrichment and confidence intervals of shared DMPs. Dots are colored according to the direction of the change in DNA methylation and shaded according to the significance of the enrichment (two-tailed Fisher’s exact test). Right. Number of DMPs within each chromatin state. **B.** Enrichment of ancestry-related DMPs in highly methylated CpGs in the genome (mean beta value >= 0.9). Left. Enrichment and confidence intervals of each tissue DMPs in highly methylated CpGs. Dots are colored according to the direction of the change in DNA methylation and shaded according to the significance of the enrichment (two-tailed Fisher’s exact test). Right. Number of DMPs within each tissue in highly methylated CpGs. **C.** Percentage of DMPs with an associated mQTL in each tissue (CpG with an identified cis-mQTL in previous study). Refer to panel B for the legend of tissue colors. **D.** Percentage of DMPs with an associated mQTL that is *cis*-driven (DMP explained by cis-genetic effects). Refer to panel B for the legend of tissue colors. **E.** Percentage of DVPs with an associated mQTL in each tissue. Refer to panel B for the legend of tissue colors. **F.** Percentage of DVPs annotated as mCpGs in which the ancestry differences are driven by previously reported *cis*-driven effects. Refer to panel B for the legend of tissue colors. **G.** Number of tissues in which the ancestry-DMPs are differentially methylated (left) and the proportion of shared ancestry-DMPs (right). High sharing: >= 4 tissues; Moderate sharing: 2–3 tissues. **H.** Number of tissues for which a CpG is differentially methylated, comparing Ancestry-DMPs that are *cis*-driven (genetically driven by an mQTL), Ancestry-DMPs that are not *cis*-driven and Ancestry-DMPs without a defined mQTL. **I.** Concordance of whole-blood ancestry-DMPs between chromatin states annotated in a European individual and an African-American individual. **J.** Example of highly shared ancestry-DMPs (CpGs differentially methylated across 8 tissues).

### The female genome is hypermethylated at Polycomb binding sites in tissue-shared DMPs

Hypermethylation in females has been previously observed in whole blood(Grant et al. 2022). Here, we wanted to address whether autosomal DNA methylation is biased toward hypermethylation in females across tissues. In all tissues, we find significantly more hypermethylated sex-DMPs in females compared to males (two-tailed binomial test; 3.22e-18 < FDR < 8.77e-240 except breast and whole blood (two-tailed binomial test; FDR > 0.05), the two most unbalanced tissues, **Fig. 4A**). Consistent with this finding, 75% of the shared sex-DMPs in more than one tissue (tissue-shared sex-DMPs, 15% of the total sex-DMPs), are significantly more methylated in females (two-tailed binomial test; p value = 9.26e-65; **Fig. 4B**).

We then wanted to determine whether hypermethylated sex-DMPs in females are associated with genes with specific functions. However, we found no biologically meaningful pathway enriched (**Table S6**). Finally, to assess replication, we compared our findings with those of an independent study in whole blood(Grant et al. 2022) (see Methods). Similarly, 74% of their sex-DMPs were hypermethylated in females, and when comparing their hypermethylated sex-DMPs with our results, we found a significant overlap (two-tailed Fisher’s exact test, p value = 1.29e-9, OR = 331).

Then, we investigated whether hypermethylated positions in females were located in particular chromatin regions (see Methods). Sex-associated changes in DNA methylation show tissue-specific enrichment in chromatin states (**Fig. 4C, Fig. S4A**), with the strongest signal found in Polycomb target regions in both sexes (bivalent TSSs in the lung and Polycomb-repressed regions in the colon) and in enhancers in the muscle and kidney. Interestingly, hypermethylated sex-DMPs in females shared in more than one tissue are preferentially located at repressed Polycomb regions, and to a lesser extent at TSS (**Fig. 4D**). Furthermore, when we evaluated the preferential location of tissue-shared sex-DMPs, including those in previous studies(Grant et al. 2022) (tissue-shared sex-DMPs shared in 2 or more tissues), we observed a higher enrichment and more significant p-value at repressed polycomb regions (OR 2.66 vs. OR 2.71; FDR 3.17e-8 vs. FDR 9.43e-9) for female-hypermethylated DMPs. To expand these results genome-wide, we evaluated the preferential location of sex-DMRs (differentially methylated regions) recovered using lung WGBS (whole-genome bisulfite sequencing) data from a previous study(Rizzardi et al. 2021) (see Methods). Similar to our array results, hypermethylated sex-DMRs in females are preferentially located at repressed Polycomb regions and TSSs (**Fig. S4B**). Finally, to further assess the sex specificity of shared sex-DMPs, we compared the DNA methylation values of tissue-shared sex-DMPs between sex-specific tissues (ovary, prostate, and testis). We observed consistent replication of the female-hypermethylation bias, with DNA methylation values being significantly higher in the ovary than in both the prostate and testis (**Fig. S4C**).

To further explore this pattern of tissue-shared hypermethylation in repressed polycomb regions, we examined whether the autosomal female hypermethylated positions were enriched at specific transcription factor-binding sites (TFBSs) based on TF ChIP-seq data from the same tissue types(Oki et al. 2018) (see Methods). We found enrichment of 30 TFBS in female hypermethylated sex-DMPs (**Table S7**). The tissue-shared female hypermethylated sex-DMPs are enriched in 177 TFBSs (**Fig. 4E, Table S7**), including TFBSs from the polycomb repressive complex (*RNF2, RING*) and other TFs involved in the maintenance of X chromosome inactivation (*ASH1L, SETDB1, SAFB*)(Migeon 2019; Chu et al. 2015; Pullirsch et al. 2010). Additionally, the 177 TFBs significantly located in shared hypermethylated sex-DMPs are enriched in the polycomb repressive complex (**Table S8**). In line with these results, multiple Polycomb repressive complex genes were consistently overexpressed in females across tissues, such as *BMI-1*, which is consistently overexpressed in females across seven different tissues (pancreatic, breast, adipose, skin, thyroid, and muscle)(García-Pérez et al. 2022). Finally, the polycomb repressive complex is responsible for the deposition of the H3K27me3 chromatin mark. Therefore, we evaluated whether sex differences in the DNA methylation of Polycomb target regions could also be detected in H3K27me3 patterns across tissues, as demonstrated in human and mouse placentas(Nugent et al. 2018). We analyzed publicly available H3K27me3 ChIP-Seq datasets from the colon sigmoid (two males and one female, see Methods) and found a strong bias toward female-specific peaks compared with male-specific peaks (324 female-biased peaks vs. 170 male-biased peaks, two-tailed binomial p value 3.9e-17). However, we did not observe this sex bias when analyzing ChIP-Seq data from other tissues, such as the heart or colon transverse.

Together, these results suggest that the female genome is consistently hypermethylated across tissues, especially at binding sites of the polycomb repressive complex.

**Figure 4.**
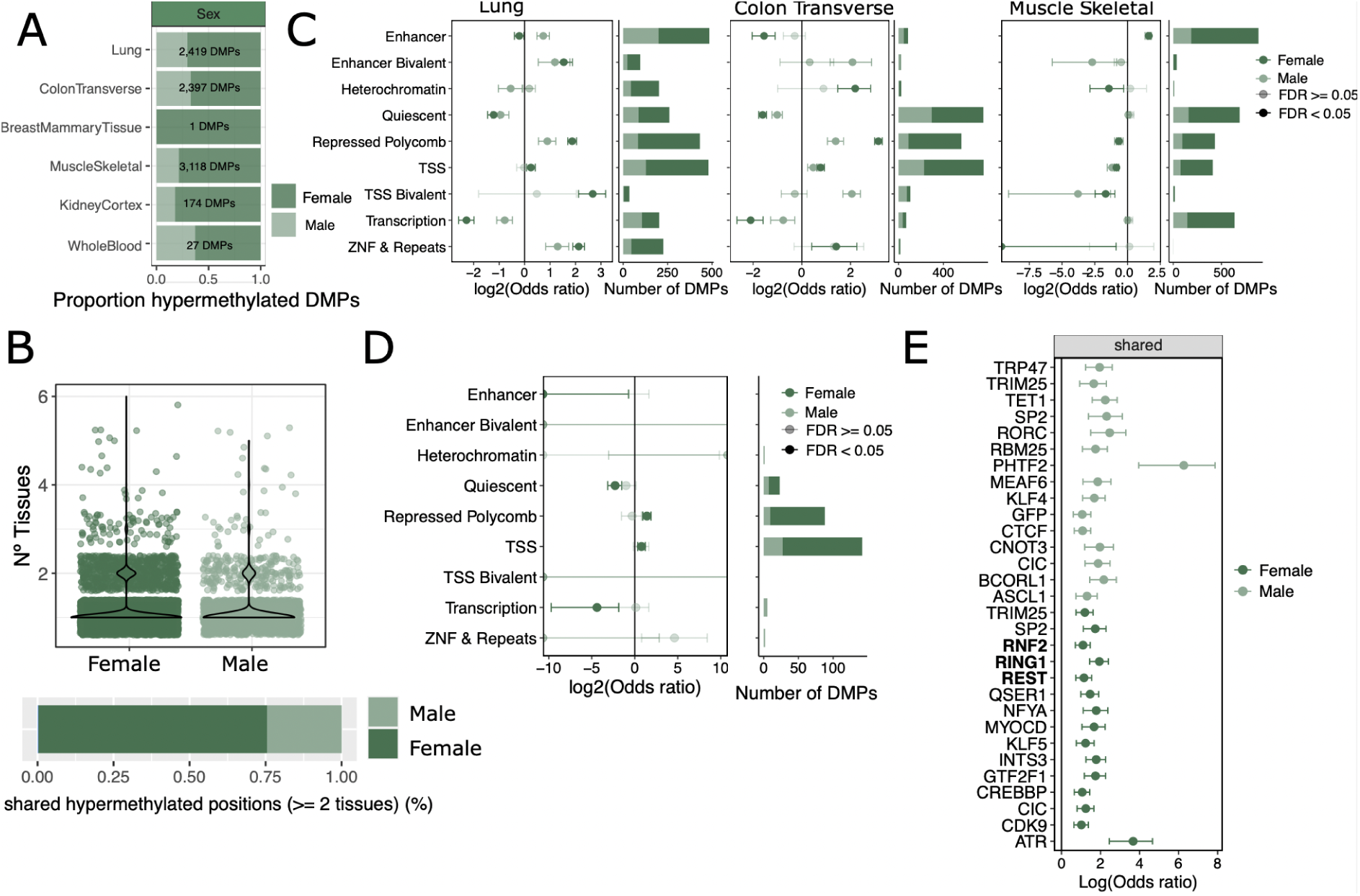
Female hypermethylation is enriched at Polycomb binding sites. **A.** Proportion of hypermethylated DMPs in males/females across tissues. Numbers represent the total amount of sex-DMPs per tissue. **B.** Number of tissues in which the sex-DMPs are differentially methylated (top) and the proportion of shared sex-DMPs in more than 1 tissue by the direction of change (bottom). **C.** Sex-DMPs enrichment at different chromatin states across the tissues with the higher number of DMPs. Left. Enrichment and confidence intervals of each tissue-DMPs. Dots are colored according to the direction of the change in DNA methylation and shaded according to the significance of the enrichment (two-tailed Fisher’s exact test). Male represents hypermethylation in males, Female represents hypermethylation in females. Right. Number of DMPs within each chromatin state per tissue. **D.** Enrichment of shared sex-DMPs (CpGs differentially methylated with sex across 2 or more tissues) in chromatin states. Left. Enrichment and confidence intervals of shared-DMPs. Dots are colored according to the direction of the change in DNA methylation and shaded according to the significance of the enrichment (two-tailed Fisher’s exact test). Male represents hypermethylation in males, Female represents hypermethylation in females. **E.** Transcription factor binding sites (TFBS) enriched at shared female hypermethylated sex-DMPs. Polycomb TFs are shown in bold.

### DNA methylation does not consistently increase with age near developmental genes in the gonads

DNA methylation changes with age have been extensively studied in different tissues(Rakyan et al. 2010; Yuan et al. 2015; Seale et al. 2022; Jain et al. 2024). However, differences in age-related methylation changes between males and females have not been explored before in other tissues beyond whole blood(Yusipov et al. 2020; McCartney et al. 2019; Teschendorff et al. 2009, 2010). To address this, we identified age-DMPs per tissue for each sex separately using linear fixed-effects models (see Methods). In most tissues, age had a greater impact on males (**Fig. S5A**), with the largest number of male-age-DMPs and female-age-DMPs located in the colon and ovary, respectively. Downsampling tissues to the same number of samples per tissue and sex shows that colon and ovary remain the tissues with the highest number of sex-age-DMPs, but we lose the ability to detect DMPs in the tissues with lower sample sizes. (**Fig. S5B**).

Next, we overlapped sex-age-DMPs with the different chromatin states defined by chromHMM in the corresponding tissues(Boix et al. 2021) to find common and sex-specific patterns of aging DNA methylation variation (see Methods). Hypermethylated age-DMPs were located across most tissues and for both sexes in chromatin states defined by the H3K27me3 mark (Polycomb target regions, i.e., bivalent TSS, bivalent enhancer, and repressed Polycomb states) (two-tailed Fisher’s exact test, FDR < 0.05) (**Fig. 5A-B**), as previously reported in other human(Teschendorff et al. 2010) and mammalian tissues(Lu et al. 2023). However, the ovaries and testes displayed methylation patterns different from those of the other tissues, with hypermethylation located in the enhancer and quiescent regions, respectively, but not in regions marked by H3K27me3 (**Fig. S5C**).

To further explore the common pattern of aging in both sexes, we focused on the overall aging-related DMPs found in the non-sex stratified age‒differential methylation analysis (**Fig. 2A**). We investigated whether the hypermethylated positions in older individuals were enriched at specific TFBSs based on TF ChIP-seq data from the same tissue types(Oki et al. 2018) (see Methods). Consistent with our previous observations, we found that, among others, TFBS that are part of the polycomb repressive complex (e.g., *CBX8*, *RNF2, EZH2, and KDM2B*) were enriched among hypermethylated age-DMPs of the colon, lung, and prostate (**Fig. 5C, Fig. S5D**) but not in the ovary or testis (**Table S9**). Finally, as the Polycomb target regions were hypermethylated in both old (**Fig. 5A-B**) and female (**Fig. 4D**) individuals, we assessed whether age- and sex-related DMPs had independent (additive) contributions to the same CpGs (see Methods). We find that the additive contribution of age and sex in colon and lung is higher than expected by chance (two-tailed Fisher’s exact test, FDR <0.05). As expected, hypermethylated CpGs in females and older individuals were significantly enriched in target regions of the polycomb repressive complex (**Fig. 5D, Fig. S5E**). Our results show that epigenetic aging in polycomb-repressed regions reflects both shared and distinct effects of sex and age. Age-related hypermethylation appears driven by DNA damage and repair, while sex differences likely stem from sustained polycomb-mediated repression of the X chromosome in females, highlighting the complex interplay between aging and sex in shaping the epigenetic landscape.

We next sought to characterize the functional pathways affected by age-methylation changes. Aging influenced the DNA methylation of genes belonging to a wide range of biological pathways, mainly from hypermethylated DMPs with age (**Fig. 5E, Table S10**), and developmental pathways being altered across four tissues (**Fig. S5F**). In addition to aging affecting developmental pathways, it also influences angiogenesis in the ovary (**Table S10**), increasing its DNA methylation with age and probably altering the expression of the associated genes (**Table S2**). Finally, to explore the relationship between changes in biological pathways and genomic location, we stratified DNA methylation changes by chromatin region. We observed that hypermethylated age-related DMPs in promoters and repressed polycomb and bivalent enhancers were located near genes involved in developmental and metabolic pathways in the lung and colon (**Table S10**).

Age-related DMPs shared across tissues may be significant contributors to phenotypic differences between individuals. Most age-related DMPs (85%) were differentially methylated in a single tissue (**Fig. 5F**). However, tissue-shared age-DMPs showed a clear bias toward hypermethylation in older individuals, with ∼84% of the shared DMPs (363 out of 435 DMPs in 4 or more tissues) consistently increasing in methylation with age (two-sided binomial test; p value=2.795349e-30), consistent with previous literature(Teschendorff et al. 2013). Tissue-shared hypermethylated age-DMPs are enriched in developmental and differentiation pathways (**Table S11**) and are preferentially located in core promoter regions (**Fig. 5G**), possibly impacting the expression of the associated genes.

Overall, we find that aging leads to the hypermethylation of developmental genes in tissues beyond whole blood, with most hypermethylation occurring at promoters and target regions of the polycomb repressive complex for both sexes, except for the ovary and testis, probably indicating reproductive-specific aging effects on DNA methylation.

**Figure 5.**
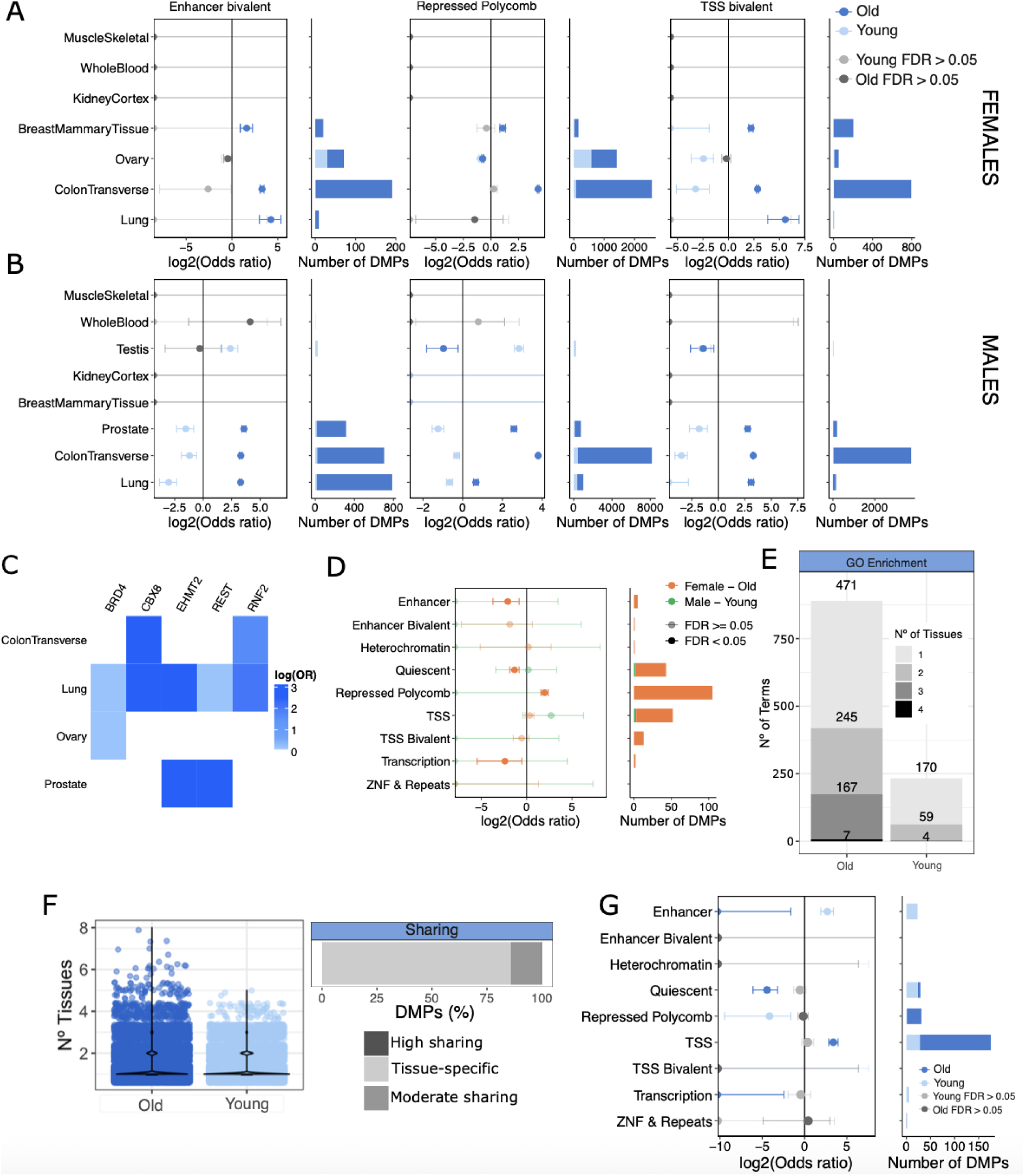
Age-associated changes in DNA methylation do not consistently target developmental genes in human gonads. **A.** Female age-related DMP enrichment at 3 chromatin states across tissues. **Left.** Enrichment of each tissue-DMPs. Dots are colored according to the direction of the change in DNA methylation. Old represents hypermethylation in older females, Young represents hypermethylation in younger females. Non-significant enrichments are colored in gray tones (two-tailed Fisher’s exact test). Confidence interval is shown. **Right.** Number of DMPs within each chromatin state per tissue. **B.** Male age-related DMP enrichment at 3 chromatin states across tissues. **Left.** Enrichment of each tissue-DMPs. Dots are colored according to the direction of the DNA methylation change and significance (two-tailed Fisher’s exact test). Old represents hypermethylation in older males, Young represents hypermethylation in younger males. Confidence interval is shown. **Right.** Number of DMPs within each chromatin state per tissue. **C.** All TFBSs enriched in hypermethylated age-DMPs that are shared between 2 tissues. OR = odds ratio. **D.** Enrichment of sex‒age additive effects across chromatin states in colon transverse. **Left.** Enrichment of DMPs. Dots are colored according to the direction of the DNA methylation change and significance (two-tailed Fisher’s exact test). Confidence interval is shown. **Right.** Number of DMPs within each chromatin state per tissue. **E.** Number of tissue-specific and tissue-shared gene ontology enriched terms for age-hyper and age-hypo-methylated positions (FDR < 0.05, GO biological process). **F.** Number of tissues in which the age-DMPs are differentially methylated (top) and the proportion of shared age-DMPs (bottom). **G.** Enrichment of shared age-related DMPs (CpGs differentially methylated with age across 3 or more tissues) in chromatin states. Dots are colored according to the direction of the change in DNA methylation. Non-significant enrichments are colored in gray tones (two-tailed Fisher’s exact test). Confidence interval is shown.

## DISCUSSION

The study of interindividual DNA methylation variation is essential for advancing our understanding of human biology and disease. Here, we jointly quantify the associations of four individual traits with DNA methylation across multiple tissues to compare their relative contributions to DNA methylation variation. We find that genetic ancestry and age are the main contributors to DNA methylation variation across tissues, with aging effects being less strong and, hence, more difficult to detect with lower sample sizes. We observe that age-, sex-, and ancestry-related DNA methylation changes at single CpGs rarely mediate gene expression differences. This likely reflects both the more stable nature of DNA methylation compared to the noisier RNA assays, and the genomic context of the affected CpGs, often in regions where methylation reinforces pre-existing expression levels rather than altering them(Ehrlich 2019; Csankovszki et al. 2001; Schlesinger et al. 2007). Moreover, previous studies suggest that such epigenetic differences may instead prime genes for differential responses under stimuli or stress, highlighting a potential regulatory role that is not apparent in healthy conditions(Netea et al. 2020; Das et al. 2019; Johnston et al. 2024).

Differences in DNA methylation between human populations have been described in whole blood(Husquin et al. 2018; Galanter et al. 2017; Giri et al. 2017; Bell et al. 2011). Here, we report a generalized difference in the DNA methylation levels of quiescent and promoter regions between African and European populations that is not influenced by differences in chromatin states. We hypothesize that the higher genetic variability in African populations(2015a) is coupled with higher DNA methylation variability in this population. This variability would be particularly evident in genomic regions with extreme methylation levels, such as highly methylated quiescent regions and unmethylated promoters, where subtle differences in methylation are easier to detect by current arrays and sample sizes. Importantly, we show that ancestry-related DNA methylation variation in promoter regions has a modest, but significant, influence on the gene expression of associated genes. This finding suggests that population-specific DNA methylation differences can contribute to gene expression variation and underscores the need of studying epigenetic diversity across ancestries. Finally, although a large fraction of ancestry-related DNA methylation variation appears to be genetically driven, only a small subset of DMPs and DVPs could be linked to cis-mQTLs, with particularly few detected in tissues with limited sample sizes. This suggests that many cis-mQTLs remain to be identified. This suggests that many potential *cis*-mQTLs still need to be identified. Therefore, future efforts are needed to elucidate the effects of genetics on DNA methylation variation in larger and more diverse cohorts and under multiple conditions and cell types.

Recent work has shown that chromatin states differ between African and European individuals(Aracena et al. 2024). Our findings build on this by demonstrating that population-specific DNA methylation differences significantly affect chromatin states that are differentially annotated between populations, suggesting a poor generalization of current chromatin annotations between human populations. Our results suggest that regulatory element location differs between populations. Considering that most studies often used European-descent samples to annotate regulatory elements(Breeze et al. 2022), these findings raise the question whether current regulatory element annotation may be biased toward European populations. Indeed, recent work has shown that current gene annotations are systematically biased towards European-descent transcripts(Clavell-Revelles et al. 2025). Expanding epigenetic profiling to underrepresented populations will therefore be essential to generate more accurate and broadly applicable regulatory maps. In addition, the differences in DNA methylation between these populations in enhancers impacting inflammation-related pathways could be linked, among other factors, to the increased levels of inflammation in African American individuals previously reported(Aracena et al. 2024; Pennington et al. 2009; Quach et al. 2016).

Hypermethylation in females has been previously observed in whole blood(Grant et al. 2022). Here, we provide evidence that the female autosomal genome is hypermethylated across tissues, specifically at Polycomb-repressed regions, and to a lesser extent at TSS. The polycomb repressive complex is known to play a role in repressing one of the female X chromosomes in the process of X chromosome inactivation. This process involves the deposition of the H3K27me3 chromatin mark throughout the X chromosome, with the last step involving the recruitment of DNA methyltransferases to methylate CpG islands(Augui et al. 2011; Gendrel et al. 2012). In line with this, multiple genes from the polycomb repressive complex (*BMI1, EZH2, SUZ12*) and genes known to interact with *XIST* (*ERH, RNMT, TRA2B*), a major effector of the X inactivation process(Migeon 2019), are upregulated in females in multiple tissues(García-Pérez et al. 2022). Importantly, in autosomes, deposition of the H3K27me3 chromatin mark has been previously reported to be female-biased in liver(Lau-Corona et al. 2020)the placenta(Nugent et al. 2018; Lau-Corona et al. 2020) impacting expression of nearby genes(Nugent et al. 2018; Lau-Corona et al. 2020). In parallel, genes that are targets of Polycomb repressive complex 2 (*PRC2*) and have the H3K27me3 chromatin mark deposition are specifically upregulated in females(Oliva et al. 2020). Thus, these findings together with our findings on higher methylation at polycomb binding sites in females could be explained by an increased activity of the polycomb repressive complex and associated DNA methyltransferases in females that extends beyond the X chromosome. Additional studies are needed to elucidate the functional implications of the hypermethylation of the female genome, specifically at Polycomb target regions

Finally, the accumulation of DNA methylation changes with age has been extensively studied(Jain et al. 2024). In particular, aging drives hypermethylation in target regions of the polycomb repressive complex across tissues(Lu et al. 2023; Teschendorff et al. 2010) and this pattern is conserved across mammalian species(Lu et al. 2023; Teschendorff et al. 2010). However, the differences in age-related methylation changes between males and females have not yet been well characterized. Here, we show that systematic hypermethylation of polycomb-target regions and regulatory regions associated with developmental genes observed previously(Teschendorff et al. 2010) occurs in both sexes. However, we did not observe the same trend in the gonads (ovary and testis). Previous findings suggest that pervasive DNA damage with aging leads to the hypermethylation of CpG islands specifically at targets of the polycomb repressive complex(O’Hagan 2014; Seale et al. 2022). Importantly, protection against DNA damage has been described in the ovary to preserve the appropriate reproductive function(Rodríguez-Nuevo et al. 2022). Additionally, the testis show high sensitivity to DNA damage, resulting in the early elimination of germ cells that acquire DNA damage with the ultimate purpose of protecting the integrity of gamete genomes(Lu and Yamashita 2017). Therefore, our findings suggest distinct aging DNA methylation trajectories in the gonads(Oakes et al. 2007; Knight et al. 2024; Uysal and Ozturk 2020; Sukur et al. 2023) likely linked to protective reproductive health to preserve gamete quality and fertility.

Overall, our multi-individual and multi-tissue approach provides an extensive characterization of the main drivers of human DNA methylation variation across individuals in healthy conditions. By establishing a baseline, our study enables future studies to distinguish disease-associated methylation changes from healthy inter-individual variation, thereby improving the identification of epigenetic biomarkers and supporting the development of more precise diagnostic and therapeutic strategies.

### Limitations of the study

Although our study employs a multi-tissue and multi-individual approach, it has certain inherent limitations that should be considered. First, the GTEx dataset has a limited representation of human populations (restricted to African Americans and European Americans) and a skewed age distribution that favors older individuals. Our findings emphasize the pressing need to incorporate individuals with diverse ancestries, as well as developmental and pediatric samples, into multi-omic analyses, as these individual traits significantly influence DNA methylation and gene expression variation. Second, our ability to detect DNA methylation changes and correlations with gene expression is constrained by reduced sample sizes and the use of arrays instead of whole-genome bisulfite sequencing, which limits statistical power and measures a subset of CpGs. Third, although we controlled for cell-type heterogeneity using PEER factors, the increasing availability of single-cell and tissue-specific methylation references will enable more refined deconvolution approaches in the future. Finally, the limited number of available tissues and the analysis of bulk samples hamper our ability to detect cell type-specific methylation changes. Taken together, future organ single-cell atlases including more tissues, donor diversity and multiple conditions will help deepen our understanding of how individual traits influence DNA methylation.

## ACKNOWLEDGEMENTS

We thank Aida Ripoll-Cladellas and Fairlie Reese for useful discussions at the end of the work. Fig. 1a was created with <u>BioRender.com</u>.

## AUTHORS’ CONTRIBUTIONS

M.M. conceived the study and designed and supervised all the analyses;

W.O. and J.M.R. analyzed the data;

W.O. led the data analysis related to differential methylation; J.M.R. led the data analysis related to differential variability.

M.M., W.O., and J.M.R. wrote the manuscript. Corresponding author

Correspondence to Marta Mele.

## AVAILABILITY OF DATA AND MATERIALS

All the GTEx protected data are available at dbGap under the accession number phs000424.v8. (https://gtexportal.org/home/protectedDataAccess). DNA methylation data is available in GEO under the accession number GSE213478. Public-access data, including QTL summary statistics and expression levels, are available on the GTEx Portal, as downloadable files and through multiple data visualizations and browsable tables (https://www.gtexportal.org), as well as in the UCSC and Ensembl browsers.

Analysis scripts are available at https://github.com/Mele-Lab/, and all the results tables derived from the analyses conducted in this paper are deposited at zenodo (doi: 10.5281/zenodo.14360407).

## ETHICS APPROVAL AND CONSENT TO PARTICIPATE

All human donors were deceased, with informed consent obtained via next-of-kin consent for the collection and banking of deidentified tissue samples for scientific research. The research protocol was reviewed by Chesapeake Research Review Inc., Roswell Park Cancer Institute’s Office of Research Subject Protection, and the institutional review board of the University of Pennsylvania.

## CONSENT FOR PUBLICATION

Not applicable.

## METHODS

### GTEx subjects

There were 838 donors (557 biological sex males and 281 biological sex females). The age of the donors ranged from 20-70 years, with most enrolled donors being older individuals. For more details on donor characteristics and sample collection, see the GTEx v8 main paper (DataS2)(The GTEx Consortium 2020). For details on the generation of the DNA methylation data and the QC pipeline, see Oliva et al.(Oliva et al. 2023), and for more details on the gene expression pipeline, see the GTEx v8 main paper(The GTEx Consortium 2020).

### Ancestry annotation

We used GTEx v8 release genotype data(The GTEx Consortium 2020) to infer ancestry for each donor. We filtered the phased GTEx v8 whole-genome sequencing variant call file (VCF) (dbGaP accession number phs000424.v8) from the reference genome Human Build 38 (hg38) with Plink 2.0 (ref) to (1) generate a pruned subset of SNPs based on Linkage disequilibrium with the option --indep-pairwise 50 10 0.1, and (2) filter variants to keep only those on chromosome 21 as a reference with the option --chr 21 and –make-bed to create a final bed file with the filtered SNPs. The final bed file was filtered to keep only individuals with European–American and African–American ancestry, as well as admixed individuals, as identified by the GTEx consortium(The GTEx Consortium 2020). We then estimated the admixture proportion for each individual based on the pruned and filtered set of SNPs using the ADMIXTURE program (v.1.3.0)(Alexander et al. 2009). ADMIXTURE was run within a two/three/four-ancestry population model in 48 threads. To define the most accurate number of ancestry populations (K parameter), we selected the model with the lowest cross-validation error (ancestry population model with K = 2).

### Differential methylation analysis

We downloaded normalized beta counts of the 754,054 CpGs from the Infinium Methylation EPIC array generated in Olivia et al.(Oliva et al. 2023) stored in GEO (GSE213478). For CpG QC, Oliva et al.(Oliva et al. 2023) excluded CpGs that had low detection P and low bead counts, cross-reactive CpGs and those overlapping a variant (or within a single base pair extension) and mapping to sex chromosomes. We used the *limma* pipeline implemented in the R package limma (v3.58.1)([CSL STYLE ERROR: reference with no printed form.]) to run linear fixed-effects models on M values(Du et al. 2010) and corrected for the following set of covariates to be as similar as possible to previously published research(Ramirez et al. 2024; García-Pérez et al. 2022):

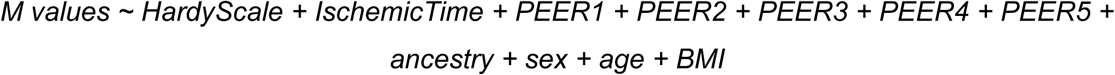

To account for hidden batch effects and other confounders of DNA methylation variance, we used the Probabilistic Estimation of Expression Residuals (PEER) method. We included 5 PEERs in the models, as it was the maximum number of PEERs available in all tissues, previously computed in Oliva et al.(Oliva et al. 2023) using inverse-normalized-transformed DNA methylation levels. These factors correct for known sources of technical variation and other confounders, such as cell type composition, as shown in the original publication(Oliva et al. 2023). We corrected all analyses for multiple testing using false discovery rate (FDR) through the Benjamini‒Hochberg method and considered CpGs differentially methylated (DMPs) at an adjusted p value below 0.05. The aging variable was categorized for visualization purposes.

To investigate the sex-specific aging effect, we divided the donors for each tissue into males and females, and we used the same *limma* pipeline and ran linear fixed-effects models on M values for each sex separately. Owing to the low variability of some of the previously included covariates, we retained only the first two PEER factors:

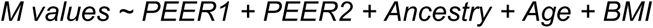

We corrected all sex-specific analyses for multiple testing separately by using false discovery rate (FDR) through the Benjamini‒Hochberg method and considered CpGs differentially methylated (sex-age-DMPs) at an adjusted p value less than 0.05.

To investigate additive effects, we conducted two-tailed Fisher’s exact tests using the fisher.test function in the R package stats (v4.3.2) with default parameters for the common DMPs between each pair of individual traits per tissue. The analysis was corrected using FDR through the Benjamini‒Hochberg method, and significant effects with adjusted p values less than 0.05 were considered. To determine whether the direction of change was concordant across individual traits, we conducted a chi-square test and corrected for multiple testing in the same way.

To validate the DMPs between ancestries, we compared our results to those obtained in an independent study using DNA methylation arrays(Aracena et al. 2024) (Table S5 from the original publication). We performed two-tailed Fisher’s exact tests to check whether the DMPs we identified overlapped with the positions observed in this previous study. The background was the set of studied positions from the EPIC array that were found in their study.

To validate the DMPs between sexes, we compared our results to those obtained with an independent dataset using DNA methylation arrays. Whole blood sex-DMPs were obtained from Additional file 1 from Grant et al.(Grant et al. 2022). We performed two-tailed Fisher’s exact tests to check whether the DMPs we identified overlapped with the positions observed in this previous study. The contingency table for Fisher’s exact test includes 4 DMPs common between both datasets, 23 DMPs only in the GTEx dataset, and 391 DMPs only in the external dataset. The background was the set of studied positions from the EPIC array that were found in their study.

### Downsampling for differential methylation analysis

To fairly compare the number of DMPs between tissues, we performed a downsampling analysis on 35 samples. We performed 50 random permutations on 35 samples from every tissue and reported the median number of DMPs per tissue and trait. We chose this approach, as downsampling to the same distribution of the tissue with the lowest sample size (skeletal muscle) would result in not enough young individuals to observe any signal in the other tissues. Using this approach also allowed us to downsample all the tissues (including skeletal muscle).

To fairly compare the number of DMPs between sexes in the sex-specific model of aging, we performed a downsampling analysis on the number of samples available for the sex with the lowest number of available donors per tissue.

### Differential variability methylation analysis

We used the function varFit from the *missMethyl([CSL STYLE ERROR: reference with no printed form.])* R package (v3.58.1) to test for differential variability running a linear fixed-effects model on M values. We corrected for the same covariates as those used in the differential methylation analysis:

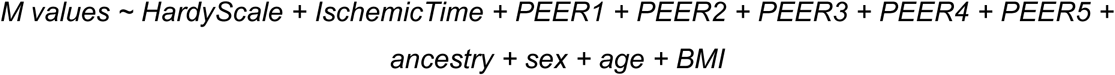

We corrected all analyses for multiple testing using false discovery rate (FDR) via the Benjamini‒Hochberg method and considered CpGs that were variably methylated at an adjusted p value below 0.05.

To investigate common differential variability in methylation and differential methylation effects, we conducted two-tailed Fisher’s exact tests on the common DMP-DVPs associated with each individual trait per tissue. The analysis was corrected using FDR through the Benjamini‒Hochberg method, and significant effects with adjusted p values less than 0.05 were considered.

We then applied the same methodology to identify differentially variable genes at the transcriptional level. We ran a linear fixed-effects model on the gene expression counts, correcting for technical covariates to have the most similar model to DNA methylation as possible:

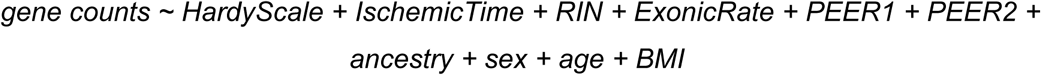

To account for hidden batch effects and other confounders of gene expression variance, we used the probability estimation of expression residuals (PEER) method. We included 2 PEERs in the models as they were mostly correlated with cell type heterogeneity (see Figure S4A from (Kim-Hellmuth et al. 2020)), and the second PEER factor was mostly correlated with the sequencing batch (see Figure S8A from (2015b)). We corrected all analyses for multiple testing using false discovery rate (FDR) via the Benjamini‒Hochberg method and considered genes that were variably expressed at an adjusted p value below 0.05.

### Annotation of DMPs

#### General classification

The DMPs were classified based on their genomic location at promoters, enhancers, gene bodies, or intergenic regions on the basis of the annotations provided in the EPIC v1.0 array manifest b5. Positions annotated as “Promoter_Associated” under “Regulatory_Feature_Group” or under “TSS200” or “TSS1500” under “UCSC_RefGene_group” were assigned as promoters. The ones with any value in “Phantom5_Enhancers” were assigned as enhancers. From the rest, the positions including “Body”, “1stExon”, “ExonBnd”, “3UTR” or “5UTR” under “UCSC_RefGene_group”, were annotated as gene bodies. The rest were annotated as intergenic. To study the enrichment of individual trait-DMPs in the different genomic locations, we performed a two-tailed Fisher’s exact test for each region separately for hypomethylated and hypermethylated DMPs. We used as background the set of studied positions from the Illumina EPIC array. The analysis was corrected using FDR through the Benjamini‒Hochberg method, and significant effects with adjusted p values less than 0.05 were considered.

#### Tissue-specific chromatin state classification

To annotate the chromatin states around each CpG, we used the 18 chromatin states inferred with ChromHMM(Ernst and Kellis 2012) generated by the ROADMAP Epigenomics consortium(Roadmap Epigenomics Consortium et al. 2015) and analyzed by EpiMap(Boix et al. 2021) for the lung (BSS01190), colon transverse (BSS01848), breast (BSS00145), testis (BSS01718), prostate (BSS01459), ovary (BSS01399), muscle skeletal (BSS01577), and kidney cortex (BSS01096) and PBMCs (BSS01419). TssFlnkD, TssFlnk, TssA and TssFlnkU are reported together under “TSS’’. EnhA1, EnhA2, EnhG1, EnhG2, and EnhWk are reported together under “Enh”. Tx and TxWk are reported together under “Tx”, and ReprPC and ReprPCWk are reported together under “ReprPC”. (**Supplementary** Fig. 3A). The data were downloaded from https://personal.broadinstitute.org/cboix/epimap/ChromHMM/observed_aux_18_hg19/CALLS/. Whole-blood chromatin states from African American and European individuals were downloaded from ENCODE (African individual ENCFF249LTX; European individual ENCFF158URQ). Colon chromatin states from 2 female and 2 male individuals were downloaded from ENCODE (Females BSS01848 and BSS01849; Males BSS01850 and BSS01851). Enhancers that were differentially annotated between populations were obtained from Table S5 from Aracena et al.(Aracena et al. 2024).

To perform enrichment of DMPs in the different chromatin states, we performed a two-tailed Fisher’s exact test, similar to the analysis of the different genomic locations. We used as background the set of studied positions from the Illumina EPIC array. To perform enrichment of DMRs in the different chromatin states, we performed a two-tailed Fisher’s exact test, similar to the analysis of the different genomic locations. We used as background the whole set of discovered DMRs and, as a significant group, the DMRs with 3 or more CpGs in the region.

#### Transcription factor-Binding Sites (TFBS)

To study the enrichment of TBFS around DMPs, we downloaded processed ChIP-seq data on transcription factors for the human v19 Lung, Kidney, Blood, Muscle, Breast, Digestive Tract, Prostate and a general catalog available in ChipAtlas(Oki et al. 2018). The transcription factors involved in the polycomb repressive complex available in the ChIP-seq data were *EZH1/2, SUZ12, YY1, KDM2B, REST, PCGF2, CBX2, CBX8, RNF2, and RYBP*.

#### Gene assignment

The probe‒gene pairs were retrieved first from the EPIC v1.0 manifest b5. We considered a probe to be part of a pair for every gene annotated under “UCSC_RefGene_Name”. For the probes annotated as promoters or enhancers for which no assigned genes were present, we annotated the closest gene via the function *matchGenes* from the R package *bumphunter* (v1.44.0)(Jaffe et al. 2012). We downloaded genes belonging to the MHC complex assembly and antigen processing pathways from the GO database (GO:0002396 and GO:0019882, respectively) for Figure 1F.

### Enrichment of Transcription Factors

We performed Fisher’s exact test for each transcription factor (TF) separately for hypomethylated and hypermethylated DMPs and adjusted for multiple testing using Benjamini‒Hochberg correction with an FDR<0.05. We used as background the TF-binding sites (TFBS) found in all tested CpGs from the Illumina EPIC array.

### Functional enrichment analyses

To perform functional enrichment analyses (overrepresentation analyses (ORA)) with the EPIC array, we need to consider that some genes contain more probes than others. For this goal, we used the *gometh* function from the *missMethyl* R package (v1.36.0)([CSL STYLE ERROR: reference with no printed form.]) to obtain the GO:BP terms. For each ORA, we carefully selected a suitable background CpG list. To investigate the biological pathways associated with each tissue- and tissue-shared DMP, we used 754,054 positions from the array as background.

### Gene expression quantification

Gene and transcript quantifications were based on the GENCODE 26 release annotation (https://www.gencodegenes.org/releases/26.html). We downloaded gene counts and TPM quantifications from the GTEx portal (https://gtexportal.org/home/datasets). We selected genes with protein-coding and lincRNA biotypes on the GTEx GENCODE v26 GTF. For the analyses involving expression data, we considered expressed genes per tissue (TPM ≥ 1 and ≥ 10 reads (unnormalized) in ≥ 20% of the tissue samples), excluding genes in the pseudoautosomal region (PAR). In total, we analyzed 22,967 genes (18,185 protein-coding genes and 4,782 lincRNAs) across tissues.

### Hierarchical partition analysis

To calculate the independent relative contributions of tissue/donor and different individual traits to DNA methylation variation, we used a variance partitioning approach using the R package variancePartition R (v1.32.5)(Hoffman and Schadt 2016). Specifically, variance partitioning decomposes the model R2 using a linear mixed model to quantify the variation in DNA methylation attributable to each variable in the model. To account for hidden batch effects and other confounders of DNA methylation variance such as cell-type composition, we used the probability estimation of expression residuals (PEER) method. We included 5 PEERs in the models, as it was the maximum number of PEERs available in all tissues, previously computed in Oliva et al.(Oliva et al. 2023) using inverse-normalized-transformed

DNA methylation levels. The model used to calculate the relative contribution of tissue and individual was:

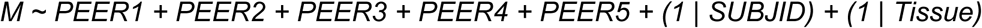

The model used to calculate the relative contribution of each individual trait was as follows:

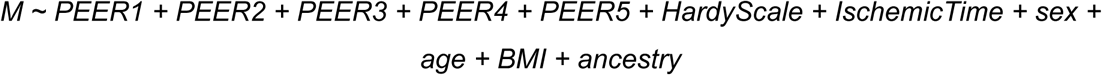

We then applied the same methodology to calculate the independent relative contribution of tissue/donor and each individual trait to the gene expression variation, using the same subset of tissues and samples. The models used are specified below:

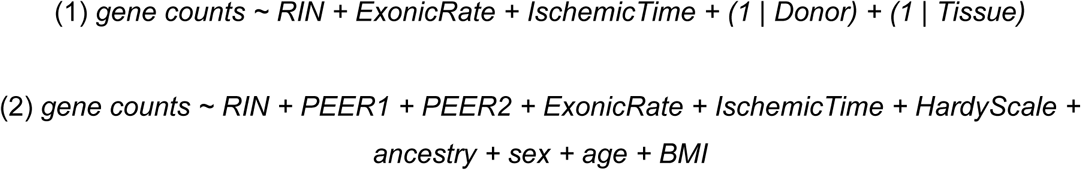

Genes or CpGs highly explained by individual were selected based on the R^2^ of individual > 0.5. Genes or CpGs highly explained by tissue were selected based on the R^2^ of tissue > 0.5.

### Identification of Cis-driven Effects

To explore whether ancestry-related DMPs/DVPs are driven by *cis* genetic effects, we focused on those previously reported as mCpGs (CpGs with significant *cis-*mQTLs)(Oliva et al. 2023). We filtered variants by a MAF > 0.01 and a window of 500 kb around the CpG. We run two different models per CpG, excluding (H0) and including (H1) the variable Ancestry while correcting in both cases for the *cis-*mQTL variants:

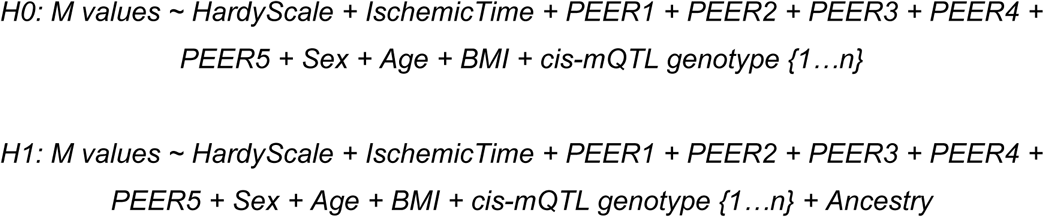

We applied the *anova* function from the R stats package (v4.3.2) to compare the models using F tests. After multiple test corrections with the Benjamini‒Hochberg method, if FDR < 0.05, we could reject the null hypothesis in which the two models are not significantly different and conclude that the ancestry effect is not solely driven by *cis* genetic effects. We considered the mCpGs associated with at least one independent *cis*-mQTL. We excluded *cis*-mQTLs if any individual was missing the genotype or if there were fewer than 3 genotyped individuals of each allelic combination. To avoid including linearly dependent *cis*-mQTLs, we computed the pairwise correlation matrix among the *cis*-mQTLs using the *findCorrelation* function in the caret R package (v6.0.94)(Kuhn 2008) and excluded highly correlated *cis*-mQTLs (correlations > 0.85).

### Enrichment of ancestry-related DMPs in *cis*-mQTLs

To investigate whether ancestry-related DMPs are overrepresented in mCpGs (CpGs with significant *cis-*mQTLs), we downloaded the latest analyses from Olivia et al.(Oliva et al. 2023) (https://gtexportal.org/home/datasets). As indicated in the GTEx portal, to obtain the list of mCpGs per tissue, we selected *cis-*mQTLs with q values < 0.05. Then, for each tissue, we computed a two-tailed Fisher’s exact test to test whether there were more ancestry-DMPs that were also mCpGs than expected if they were independent. We used as background the set of studied positions from the Illumina EPIC array. We determined a significant enrichment after correcting for multiple testing via the Benjamini‒Hochberg method across tissues at an FDR < 0.05.

### Enrichment of ancestry-related DMPs in regions with extreme methylation status

To assess whether African-DMPs were enriched in regions with extreme DNA methylation profiles, such as promoters or quiescent regions, we first generated a catalog with the coordinates of those regions. Briefly, we selected CpGs that, across donors, had average methylation levels for a specific tissue above 0.9 and CpGs that had average methylation levels below 0.1. We then computed a two-tailed Fisher’s exact test to test whether there were more ancestry-DMPs located in those regions than expected (using as background the location of all the tested CpGs from the Illumina EPIC array). We determined a significant enrichment after correcting for multiple testing via the Benjamini‒Hochberg method across tissues at an FDR < 0.05.

### Correlation between DNA methylation and gene expression

The probe‒gene pairs were retrieved first from the EPIC v1.0 manifest b5. We considered a probe to be part of a pair for every gene annotated under “UCSC_RefGene_Name”. For the probes annotated as promoters or enhancers for which no assigned genes were present, we annotated the closest gene using the function *matchGenes* from the R package *bumphunter* (v1.44.0)(Jaffe et al. 2012). The total numbers of CpGs tested in promoters, enhancers, and gene bodies are 108,810, 25,287, and 370,343, respectively. We used the DNA methylation and gene expression residuals, after regressing out the effects of cell type composition (PEER factors) and technical covariates. We computed Pearson’s correlation for the DMP-DEG pairs and corrected for multiple testing using the Benjamini‒Hochberg method separately per genomic region (enhancer, promoter, and gene-associated). We considered correlations with an FDR < 0.05 to be significant. To compute the percentage of DEGs that significantly correlated with DMPs, we considered DEGs associated with at least one probe in the array. To increase our detection power, we repeated the analysis considering as DMP-DEG pairs all the DMPs significant at a p value nominal < 0.05.

### Mediation analysis

To identify genes whose changes in expression with individual traits were mediated by the DNA methylation of the associated CpGs, we used a regularized mediation analysis based on penalized structural equation modeling implemented in the R package *regmed* (v2.1.0)(Schaid et al. 2022; Schaid and Sinnwell 2020). For each DEG for a particular tissue and individual trait, we considered all the associated DMPs as mediators. To avoid including linearly dependent CpGs, we computed the pairwise correlation matrix among the DMPs using the *findCorrelation* function in the caret R package (v6.0.94)(Kuhn 2008) and excluded highly correlated CpGs (correlations > 0.9). We followed the published pipeline to run mediation with one exposure and multiple mediators with a penalty “lambda” grid of *seq (from = 0.6, to = 0.01, by = -0.05)*. We then selected the best model with the function regmed.grid.bestfit and considered all those genes with at least one CpG having an alpha*beta different from 0 to be mediated.

### Ancestry-specific chromatin regions

To address whether Ancestry-DMPs were located at genomic regions that had different chromatin states between African and European human populations, we first created a catalog of CpGs with an annotation of whether the chromatin state of its location was the same (“Common region”) or different (“Ancestry-specific”) between these populations. Additionally, we focused on the “Ancestry-specific” regions and computed a two-tailed Fisher’s exact test to test whether there were more ancestry-DMPs located in any of those regions than expected (using as background the location of all the tested CpGs from the Illumina EPIC array), separately for hypermethylated and hypomethylated positions. We determined a significant enrichment after correcting for multiple testing via the Benjamini‒Hochberg method across tissues at an FDR < 0.05.

### Differential peak calling between sexes

To assess whether sex-related DMPs are related to differential Polycomb repressive complex binding, we analyzed publicly available ChIP-Seq data for the H3K27me3 chromatin mark in heart and colon tissue: 4 heart, 3 colon sigmoid, and 4 colon transverse samples (male samples: GSM7247417, GSM7247418, GSM6347302, GSM6347303, GSM6347304, GSM4250725, GSM4250726, GSM4146448, GSM4146449, GSM4250562, GSM4250563, GSM4250564, and GSM4250565; female samples: GSM5669434, GSM5669435, GSM5669390, GSM5669389, GSM4247242, GSM4247243, GSM5112491, GSM5112492, GSM5112493, GSM5112494, and GSM2701098). We then used the differential peak calling method from the ChIPAtlas (Diff Analysis)(Oki et al. 2018), with default parameters.

### Mapping and quality control of WGBS data

We downloaded fastq files for the WGBS data of lung GTEx samples from dbGaP under the accession number phs000424. We trimmed reads of their adapter sequences using Trim Galore (v0.6.6)([CSL STYLE ERROR: reference with no printed form.]) with a quality trimming cutoff of 30. We then aligned the trimmed reads to the hg38 build of the human genome using Bismark (v0.23.1)(Krueger and Andrews 2011) with default parameters. We then used bismark_methylation_extractor to summarize the number of reads supporting a methylated cytosine and an unmethylated cytosine for every cytosine present in the reference genome. Consistent with previous studies(Rizzardi et al. 2021), we included the parameters *“--ignore 2 --ignore_r2 3 --ignore_3prime 1 --ignore_3prime_r2 3”*. The final report file summarizes the evidence of methylation at each cytosine in the human reference genome.

### DMR analysis

We used the R package *bsseq* to analyze the WGBS data(Hansen 2024). Differentially methylated regions (DMRs) were identified as previously described(Rizzardi et al. 2021), with a minimum of 70 CpGs required in a smoothing window. Briefly, we used BSmooth(Hansen et al. 2012) to estimate CpG methylation levels. We ran a ‘small’ smooth to estimate the methylation level in a small genomic region (smoothing over windows of at least 1 kb containing at least 70 CpGs). Following smoothing, we analyzed all CpGs that had a sequencing coverage of at least 1 in all samples for a total of 16,290,182 CpGs. Finally, we computed t-statistics and identified DMRs by thresholding the t-statistics following the *bsseq* pipeline. We considered as significant DMRs those with at least 3 CpGs in them.

## List of supplementary tables

Supplementary Table 1: GO enrichment results for A) tissue-variable CpGs; B) tissue-variable CpGs per genomic location; C) individual-variable CpGs; and D) individual-variable CpGs per genomic location.

Supplementary Table 2: GO enrichment results for correlated DEGs correlated with DMPs in the ovary with age.

Supplementary Table 3: Correlation analysis between DMPs and DEGs per tissue and trait. Significantly correlated DMP‒DEG pairs are included.

Supplementary Table 4: Mediation results. Table depicting DEGs and the number of *cis*-associated DMPs that significantly mediate gene expression changes.

Supplementary Table 5: GO enrichment results for ancestry-related DMPs located in enhancers.

Supplementary Table 6: GO enrichment results for sex-related DMPs. Hyper: Higher methylation in females, Hypo: Higher methylation in males.

Supplementary Table 7: Enrichment of TFBSs in sex-DMPs A) per tissue and B) tissue-shared (DMPs shared between 2 or more tissues). Hyper: Higher methylation in females, Hypo: Higher methylation in males.

Supplementary Table 8: GO enrichment results for the TFs enriched among hypermethylated sex-DMPs shared between 2 or more tissues.

Supplementary Table 9: Enrichment of TFBSs in age-DMPs per tissue and direction of methylation change. Hyper: Increased methylation in older individuals; Hypo: Increased methylation in young individuals.

Supplementary Table 10: GO enrichment results for age-related DMPs per A) tissue and B) tissue and genomic location. Hyper: Increased methylation in older individuals; Hypo: Increased methylation in young individuals.

Supplementary Table 11: GO enrichment results for tissue-shared age-hypermethylated DMPs (more methylated in older individuals in 3 or more tissues).

## Supplementary Figures

**Figure S1.**
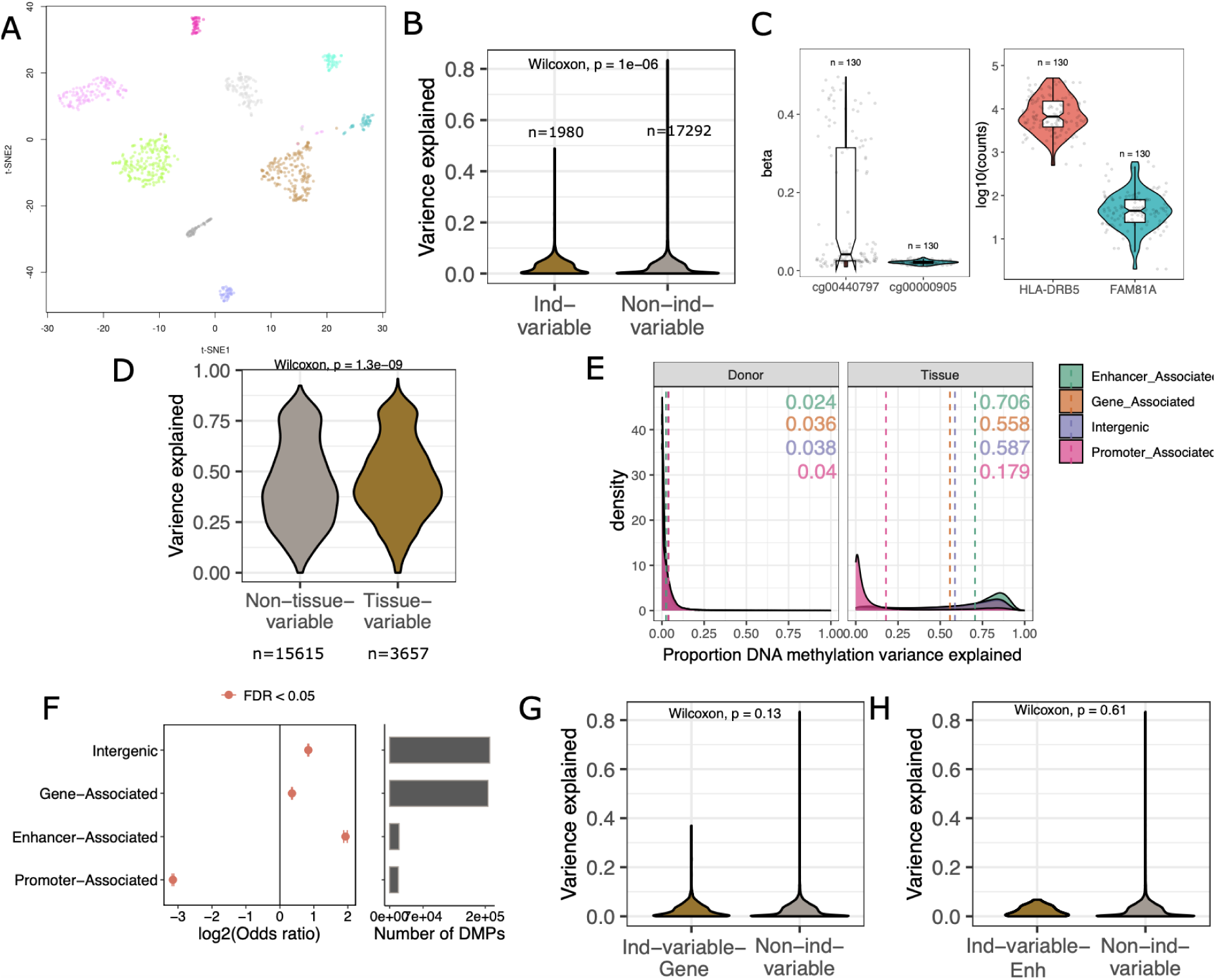
Overview of DNA methylation across tissues and individuals. **A.** t-SNE showing the clustering of the DNA methylation data by tissue. **B.** Proportion of gene expression variation explained for genes individually-variable (brown) and not individually-variable (gray) at the DNA methylation level. **C.** Examples of 1 individually-variable CpG (cg00440797) and one non-individually variable CpG (cg00000905), and their corresponding genes (HLA-DRB5 and FAM81A, respectively). **Left.** Beta values for each CpG in lung. **Right.** log10(counts) for each gene in lung. The number of individuals per group are indicated. **D**.Proportion of gene expression variation explained for genes tissue-variable (brown) and not tissue-variable (gray) at the DNA methylation level. **E.** Proportion of DNA methylation variation explained by individual (Left) and tissue (Right). The distributions are clustered by the genomic location of the CpGs. Median numbers for each genomic location are shown in each facet. **F. Left.** Enrichment of each genomic location on tissue variable CpGs. X-axis represents the log2(OR) and dots are colored according to the significance of the enrichment (two-tailed Fisher’s exact test). Confidence interval is shown. **Right.** Number of tissue variable CpGs associated with each genomic location. G. Proportion of gene expression variation explained for genes individually-variable (brown) and not individually-variable (gray) at the DNA methylation level of their gene bodies. H. Proportion of gene expression variation explained for genes individually-variable (brown) and not individually-variable (gray) at the DNA methylation level of their enhancers.

**Figure S2:**
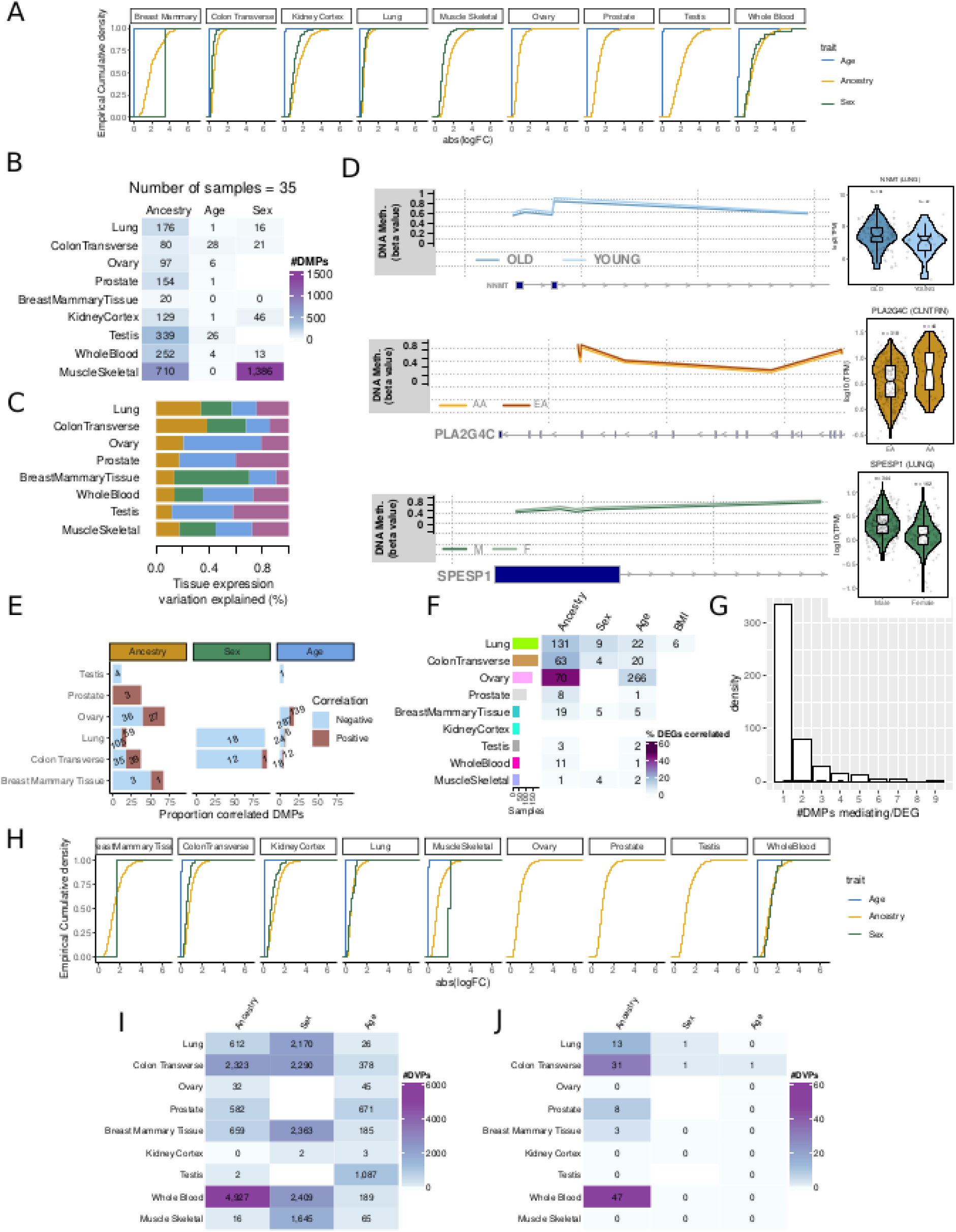
DNA methylation variation across tissues and individual traits. **A.** Effect sizes of DMPs per individual trait. **B.** Mean number of DMPs per tissue and individual trait for 50 random permutations when downsampling to an equal number of samples across tissues=35 . **C.** Proportion of total tissue gene expression variation explained by each individual trait. **D.** Examples of correlated DEGs with at least one DMP. log10(TPM) values for gene expression are shown in violin plots. Beta values for DNA methylation of correlated DMPs are plotted according to their genomic location. The gene structure is shown below each plot. **E.** Proportion and number of DMPs correlated with DEGs per tissue and individual trait., colored by the direction of correlation. **F.** Number of DEGs correlated with at least one DMP, DMPs selected based on nominal p.value < 0.05. Cells are colored by the proportion of DEGs correlated out of the total number of DEG with a probe. **G.** Number of DMPs mediating gene expression changes per each DEG. **H.** Effect sizes of DVPs per individual trait and tissue. **I.** Number of DVG per trait and tissue. **J.** Number of DVG per trait and tissue that have an associated DVP.

**Figure S3.**
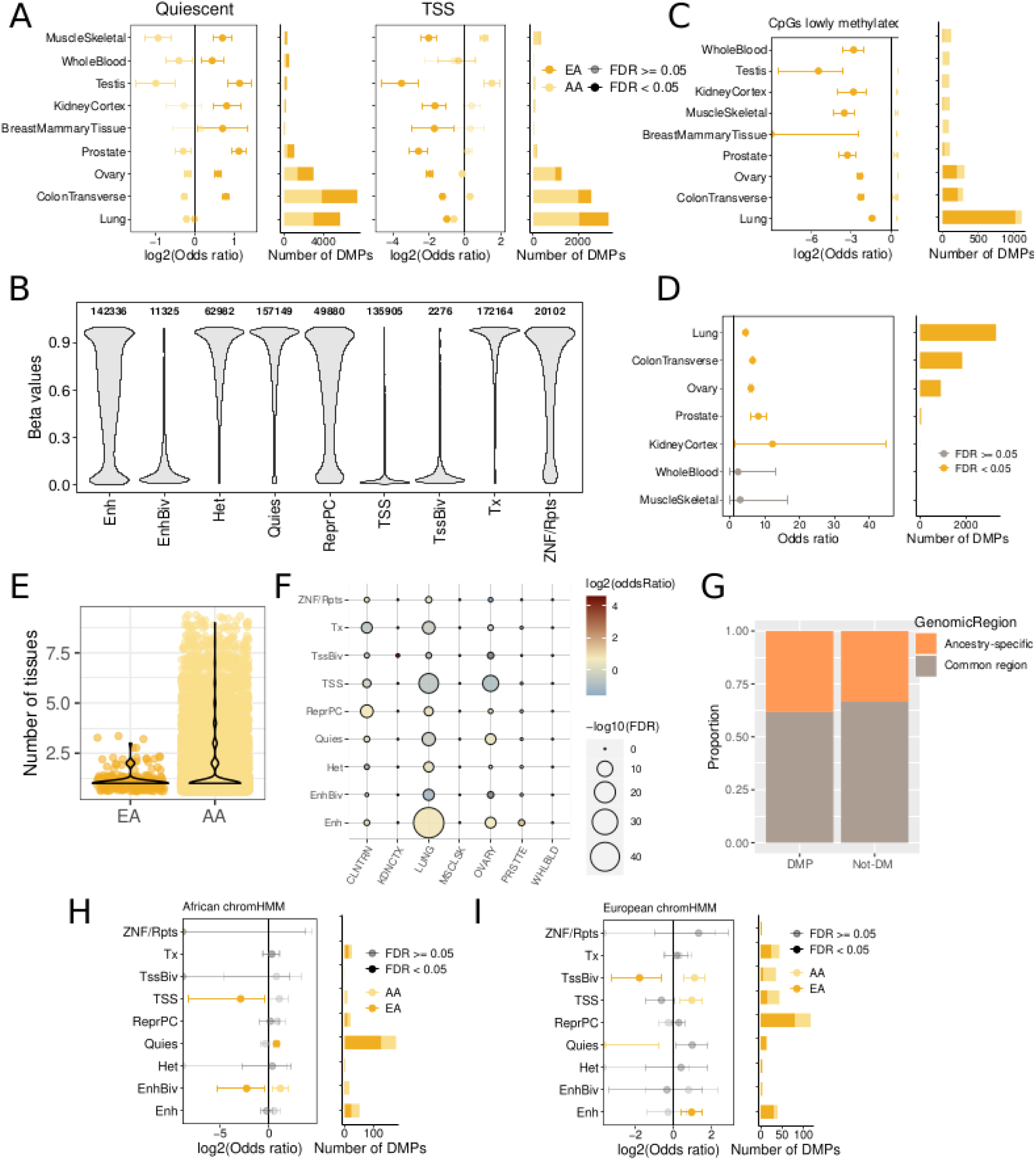
Ancestry-associated DNA methylation changes. **A.** Ancestry-DMPs enrichment at 2 chromatin states across tissues: quiescent and transcription start sites. Left. Enrichment and confidence interval of each tissue-DMPs. Dots are colored according to the direction of DNA methylation change and shaded according to the significance of the enrichment (two-tailed Fisher’s exact test). Right. Number of DMPs within each chromatin state per tissue. **B.** Representation of the different chromatin states analyzed throughout the paper and their associated DNA methylation status. **C.** Enrichment of ancestry-DMPs in CpGs not methylated in the genome. Legend is shared with panel **A**. **D.** Enrichment of Ancestry-DMPs in mQTLs across tissues. Left. Enrichment and confidence interval of each tissue-DMPs. Dots are colored according to the significance of the enrichment (two-tailed Fisher’s exact test). Right. Number of DMPs with mQTLs within each tissue. **E.** Number of tissues where ancestry-DVPs are found. **F.** Enrichment of cis-driven Ancestry-DMPs at different chromatin states across tissues. **G.** Proportion of DMPs and CpGs not DM (Not DM) overlapping Ancestry-specific genomic regions (CpGs located in a chromatin region differentially annotated between population by chromHMM). **H.** Enrichment of Ancestry-DMPs in an African-descent individual at multiple chromatin states (see Methods). **I.** Enrichment of Ancestry-DMPs in a European-descent individual at multiple chromatin states (see Methods).

**Figure S4.**
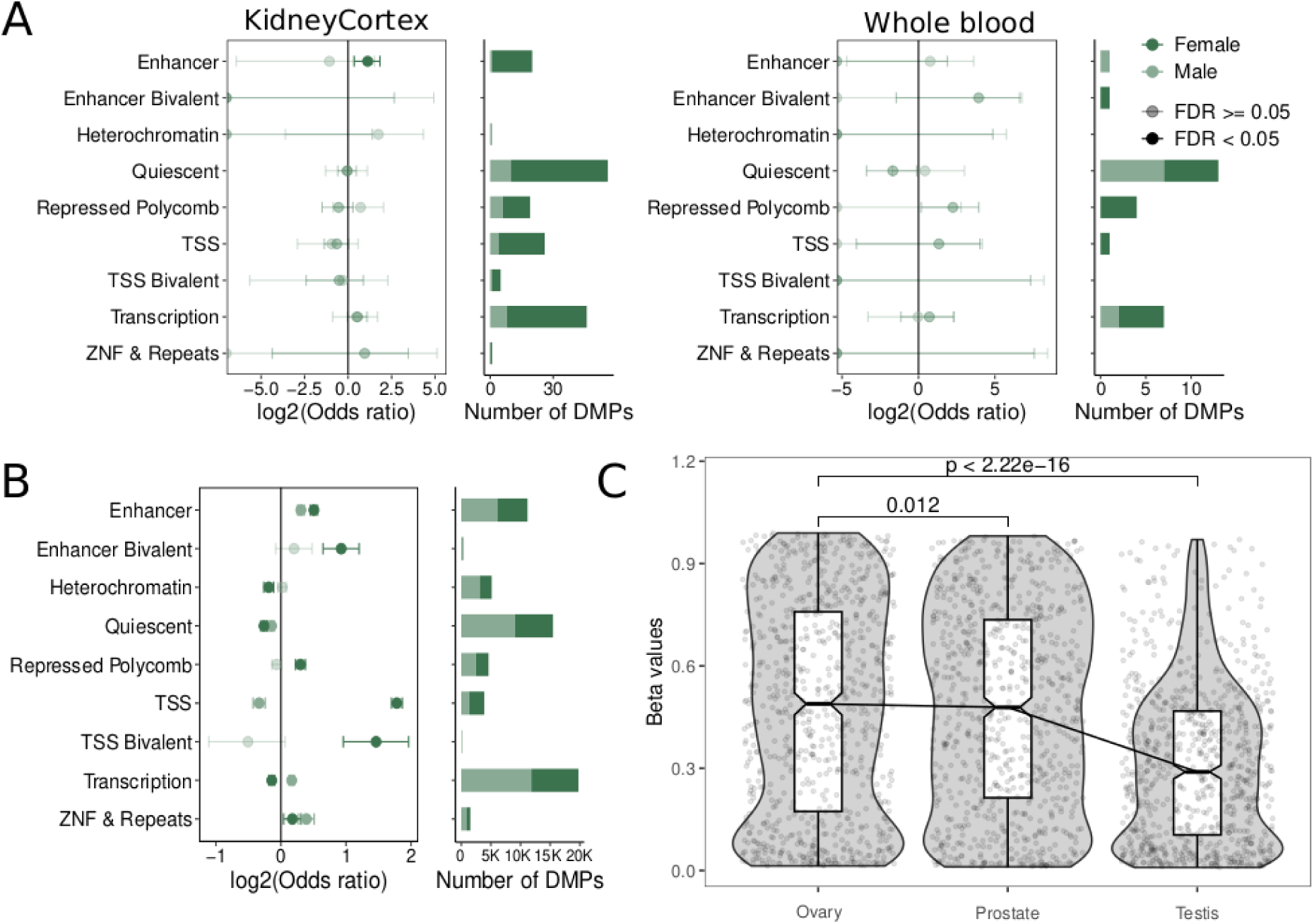
Sex-associated DNA methylation changes. **A.** Sex-DMPs enrichment at different chromatin states across tissues. Left. Enrichment and confidence interval of each tissue-DMPs. Dots are colored according to the direction of DNA methylation change and shaded according to the significance of the enrichment (two-tailed Fisher’s exact test). Right. Number of DMPs within each chromatin state per tissue. **B.** Sex-DMRs enrichment at different chromatin states in Lung. Enrichment and confidence interval of each tissue-DMPs. Dots are colored according to the direction of DNA methylation change and shaded according to the significance of the enrichment (two-tailed Fisher’s exact test). Right. Number of DMPs within each chromatin state per tissue. **C.** DNA methylation status (Beta values) of shared DMPs in the sex-related tissues. P-values correspond to Wilcoxon-test with paired data.

**Figure S5.**
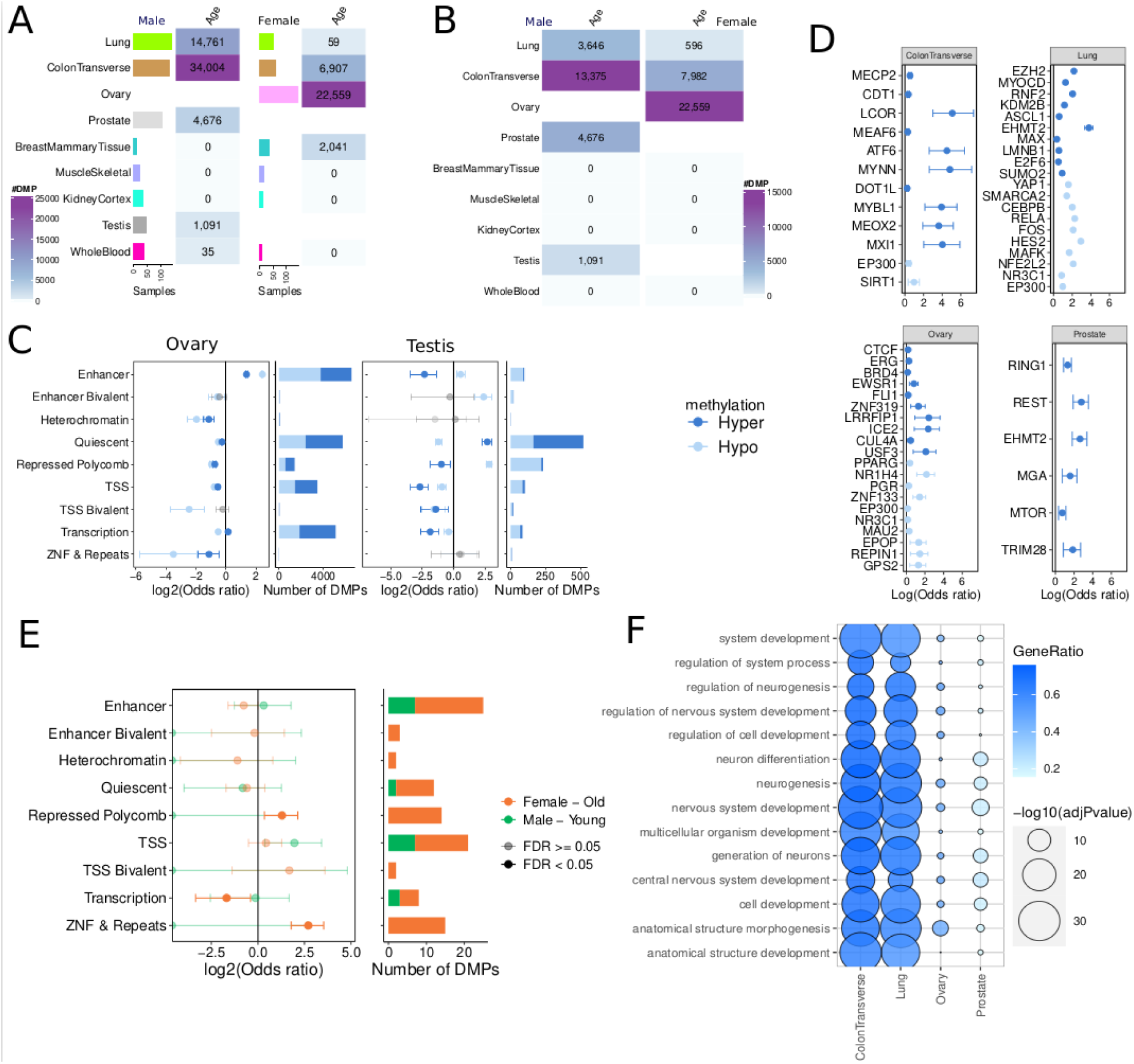
Age-associated DNA methylation changes. **A.** Number of age-DMPs per tissue in male and female individuals. Left barplot represents the number of samples for every tissue. **B.** Number of age-DMPs per tissue in male and female individuals, downsampling to the same number of individuals per sex-tissue pair. Left barplot represents the number of samples for every tissue. **C.** Age-DMPs enrichment at chromatin states for Ovary and Testis. **Left.** Enrichment OR and confidence intervals. Dots are colored according to the direction of DNA methylation change. Non-significant enrichments are colored in gray tones (two-tailed Fisher’s exact test). **Right**. Number of DMPs within each chromatin state per tissue. **D.** Top significant Transcription Factor Binding Sites (TFBS) enriched at age-DMPs for each tissue (ordered by FDR and direction of methylation change). **E.** Enrichment of sex-age additive effects across chromatin states in Lung. Left. Enrichment of DMPs. Dots are colored according to the direction of DNA methylation change and significance (two-tailed Fisher’s exact test). Confidence interval is shown. Right. Number of DMPs within each chromatin state per tissue. **F.** Enrichment for the 14 GO biological-process pathways enriched in 4 tissues.

